# De novo indels within introns contribute to ASD incidence

**DOI:** 10.1101/137471

**Authors:** Adriana Munoz, Boris Yamrom, Yoon-ha Lee, Peter Andrews, Steven Marks, Kuan-Ting Lin, Zihua Wang, Adrian R. Krainer, Robert B. Darnell, Michael Wigler, Ivan Iossifov

## Abstract

Copy number profiling and whole-exome sequencing has allowed us to make remarkable progress in our understanding of the genetics of autism over the past ten years, but there are major aspects of the genetics that are unresolved. Through whole-genome sequencing, additional types of genetic variants can be observed. These variants are abundant and to know which are functional is challenging. We have analyzed whole-genome sequencing data from 510 of the Simons Simplex Collections quad families and focused our attention on intronic variants. Within the introns of 546 high-quality autism target genes, we identified 63 de novo indels in the affected and only 37 in the unaffected siblings. The difference of 26 events is significantly larger than expected (p-val = 0.01) and using reasonable extrapolation shows that de novo intronic indels can contribute to at least 10% of simplex autism. The significance increases if we restrict to the half of the autism targets that are intolerant to damaging variants in the normal human population, which half we expect to be even more enriched for autism genes. For these 273 targets we observe 43 and 20 events in affected and unaffected siblings, respectively (p-value of 0.005). There was no significant signal in the number of de novo intronic indels in any of the control sets of genes analyzed. We see no signal from de novo substitutions in the introns of target genes.

## Introduction

We have made great strides in our understanding of the genetic determinants of autism over the past decade. These come largely from the search for new germ line (de novo) mutations in simplex families, that is, those with a single affected child. The major signal comes from exome sequence data, and in particular from the mutations that disrupt protein coding sequences [1, 2]. The best estimate of the contribution from de novo mutation derives from the observed differential incidence rates in affected and unaffected siblings, and extrapolates to about 30%. Using a variety of methods for analysis of the number of recurrent gene targets, we can further estimate that the number of strongly penetrant causal targets for de novo mutation is on the order of 500 genes [1]. Using the observation that target genes, and especially recurrent target genes, are enriched for genes under strong negative selective pressure in humans, we can now identify on the order of 200 excellent candidate target genes, those that are both targets and under strong selective pressure [3].

Potentially, we can learn more from whole genome sequencing data, although the rules for interpreting such data are not yet clear. Two recent reports that studied the relationship between non-coding variants and autism demonstrate these difficulties and the need for analysis of whole-genome data from large collations [4, 5]. In this comparatively large study, we focus on mutations within introns.

Several observations show that abnormal splicing is a major mechanism for damaging alleles. About 50% of the genetic variants underling NF1 [6] and ATM [7] result in abnormal splicing. Also, more than 50% of the variants associated with human phenotypes in the GWAS catalog [8] are within introns. With the whole genome sequencing data, we are for the first time able to systematically examine the contribution to autism from intronic mutations.

In this study, we compare the incidence of de novo mutation within the introns of affected and unaffected children from the SSC, within all genes, and within target genes. Although we see no significant differences over all genes, we find a statistically significant excess of de novo intronic indels in suspected autism target genes. We see no signal from de novo intronic substitutions. We estimate by extrapolation of the known target gene class size that de novo indels in introns of target genes contribute to about 10% of the affected within simplex families. In the Discussion, we further revise upwards our estimate of the total contribution of de novo events to autism.

## Results

### Counts and significance of intronic events

We have whole genome sequencing from 510 quad families from the Simons Simplex Collection (SSC) [9]. The first 510 families were chosen to have no de novo LGDs or CNVs in the exomes of the children. We catalogued for all de novo substitutions and indels (of size not exceeding 50 bp) using the multinomial genotyper we have previously employed [10]. All ∼2000 de novo intronic indels (DIIN) and all ∼20,000 de novo intronic substitutions (DISB) are listed in Supplementary Tables I and 2 by event, and by gene in Supplementary Table 3. We did not validate any of the DISB, as previous experience indicates that almost all would be confirmed. We validated several dozen of the DIIN using previous methods [10], and only 4% were false positives, similar to our rates from whole exome sequencing [1], and not sufficiently large to cast doubt on the findings we now describe.

The counts of de novo intronic events are summarized in Table 1. These are separated into DIIN (top half of Table 1) and DISB (bottom half of Table 1), as ‘events in affected’ or ‘events in unaffected’ siblings. The counts are for events in ‘all genes’ or divided into classes of genes by the type of target (the rows defined in column ‘gene set’), with the ‘number of genes’ in a target type as tabulated. The first sub-type is called ‘affected LGD targets’ contains the 546 genes that have been targeted by de novo LGD mutations in 5,000 affected children. We further divide the ‘affected LGD targets’ in two equal halves based on ‘protection’. Protection is the extent to which each of the genes is under purifying selection reflected by the extent of damaging mutations found in the human population [3]. The first half contains the more protected LGD targets (‘affected LGD targets, protected’) and the second half contains the less protected LGD targets (‘affected LGD targets, unprotected’). We analyzed five additional control gene sets defined based on observed de novo missense and synonymous mutation in the ∼5,000 affected children or based on observed de novo LGD, missense, and synonymous mutations in ∼2,000 unaffected children. The difference in counts of events between discordant siblings is called ‘delta’.

**Table 1.**
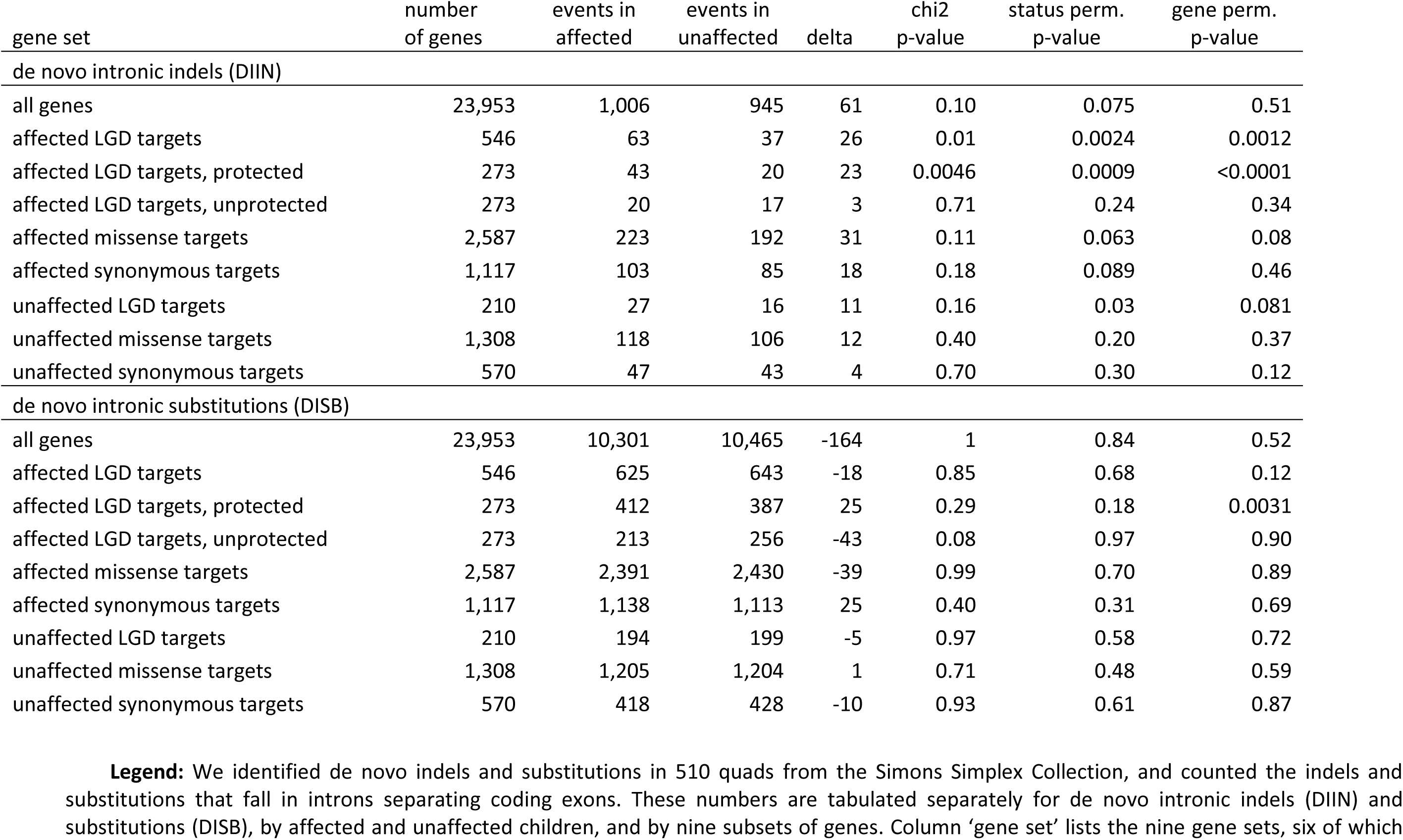
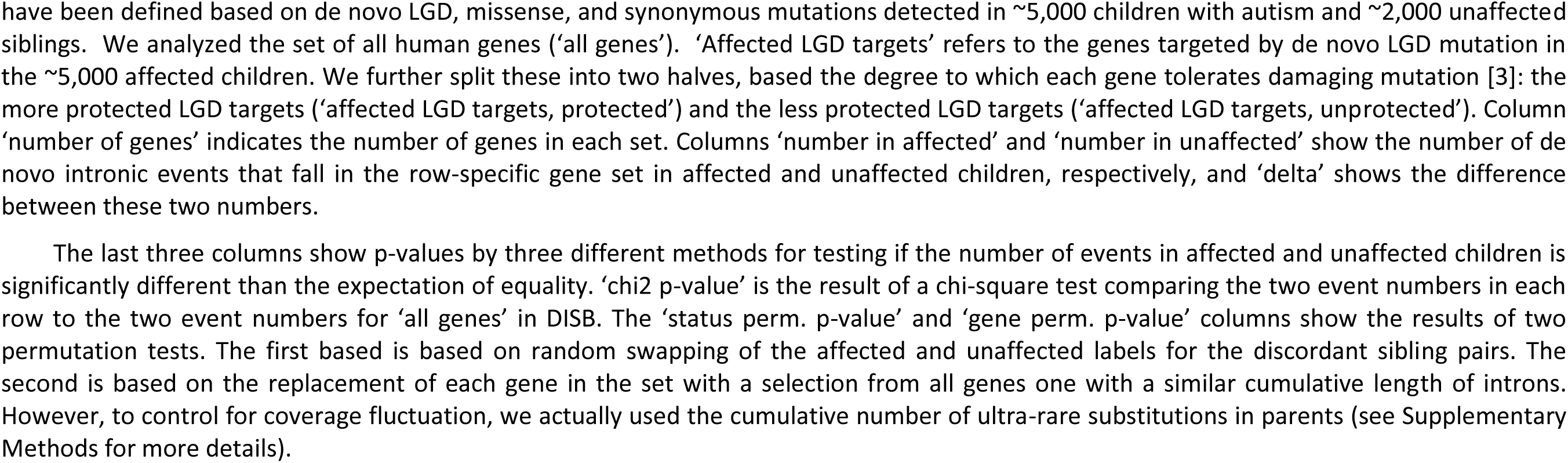
De novo intronic indels (DIIN) and substitutions (DISB) in introns between coding exons

| gene set | number<br>of genes | events in<br>affected | events in<br>unaffected | delta | chi2<br>p-value | status perm.<br>p-value | gene perm.<br>p-value |
| --- | --- | --- | --- | --- | --- | --- | --- |
| de novo intronic indels (DIIN) |  |  |  |  |  |  |  |
| all genes | 23,953 | 1,006 | 945 | 61 | 0.10 | 0.075 | 0.51 |
| affected LGD targets | 546 | 63 | 37 | 26 | 0.01 | 0.0024 | 0.0012 |
| affected LGD targets, protected | 273 | 43 | 20 | 23 | 0.0046 | 0.0009 | <0.0001 |
| affected LGD targets, unprotected | 273 | 20 | 17 | 3 | 0.71 | 0.24 | 0.34 |
| affected missense targets | 2,587 | 223 | 192 | 31 | 0.11 | 0.063 | 0.08 |
| affected synonymous targets | 1,117 | 103 | 85 | 18 | 0.18 | 0.089 | 0.46 |
| unaffected LGD targets | 210 | 27 | 16 | 11 | 0.16 | 0.03 | 0.081 |
| unaffected missense targets | 1,308 | 118 | 106 | 12 | 0.40 | 0.20 | 0.37 |
| unaffected synonymous targets | 570 | 47 | 43 | 4 | 0.70 | 0.30 | 0.12 |
| de novo intronic substitutions (DISB) |  |  |  |  |  |  |  |
| all genes | 23,953 | 10,301 | 10,465 | -164 | 1 | 0.84 | 0.52 |
| affected LGD targets | 546 | 625 | 643 | -18 | 0.85 | 0.68 | 0.12 |
| affected LGD targets, protected | 273 | 412 | 387 | 25 | 0.29 | 0.18 | 0.0031 |
| affected LGD targets, unprotected | 273 | 213 | 256 | -43 | 0.08 | 0.97 | 0.90 |
| affected missense targets | 2,587 | 2,391 | 2,430 | -39 | 0.99 | 0.70 | 0.89 |
| affected synonymous targets | 1,117 | 1,138 | 1,113 | 25 | 0.40 | 0.31 | 0.69 |
| unaffected LGD targets | 210 | 194 | 199 | -5 | 0.97 | 0.58 | 0.72 |
| unaffected missense targets | 1,308 | 1,205 | 1,204 | 1 | 0.71 | 0.48 | 0.59 |
| unaffected synonymous targets | 570 | 418 | 428 | -10 | 0.93 | 0.61 | 0.87 |

**Legend:** We identified de novo indels and substitutions in 510 quads from the Simons Simplex Collection, and counted the indels and substitutions that fall in introns separating coding exons. These numbers are tabulated separately for de novo intronic indels (DIIN) and substitutions (DISB), by affected and unaffected children, and by nine subsets of genes. Column 'gene set' lists the nine gene sets, six of which
have been defined based on de novo LGD, missense, and synonymous mutations detected in ~5,000 children with autism and ~2,000 unaffected siblings. We analyzed the set of all human genes ('all genes'). 'Affected LGD targets' refers to the genes targeted by de novo LGD mutation in the ~5,000 affected children. We further split these into two halves, based the degree to which each gene tolerates damaging mutation [3]: the more protected LGD targets ('affected LGD targets, protected') and the less protected LGD targets ('affected LGD targets, unprotected'). Column 'number of genes' indicates the number of genes in each set. Columns 'number in affected' and 'number in unaffected' show the number of de novo intronic events that fall in the row-specific gene set in affected and unaffected children, respectively, and 'delta' shows the difference between these two numbers.
The last three columns show p-values by three different methods for testing if the number of events in affected and unaffected children is significantly different than the expectation of equality. 'chi2 p-value' is the result of a chi-square test comparing the two event numbers in each row to the two event numbers for 'all genes' in DISB. The 'status perm. p-value' and 'gene perm. p-value' columns show the results of two permutation tests. The first based is based on random swapping of the affected and unaffected labels for the discordant sibling pairs. The second is based on the replacement of each gene in the set with a selection from all genes one with a similar cumulative length of introns. However, to control for coverage fluctuation, we actually used the cumulative number of ultra-rare substitutions in parents (see Supplementary Methods for more details).

The remaining columns reflect three distinct methods for determining the significance of the delta. The first method (column ‘chi2 p-value’) is based on a chi-square test. The second and third methods are based on 10,000 permutations to develop empirical distributions on delta for each row. The p-value is the proportion of permuted deltas that were greater or equal to the empirically observed delta. For the column ‘status perm. p-value’ in each permutation we randomly assign the affected and unaffected status labels of sibling pairs. In the column ‘gene perm. p-value’, we randomly select genes with similar cumulative intron length. The second and third methods are meant to guard against outlier families and outlier genes, respectively, which could give rise to spurious statistical significance in the first method. All three methods are in good agreement. See Table 1 legend and methods for additional details.

### Signal from indels in likely autism genes

The counts for DISB in all genes are 10,301 and 10,465 for affected and unaffected, respectively, with a delta of -164. Clearly, these are not significantly different. The rates average to 1.2*10^-8^ per highly covered base pair per child, a number in keeping with previous rates for de novo mutation over the whole-genomes [11-16]. The counts for DIIN in all genes are 1006 and 945, with a delta of 61, also without statistical significance (Table 1). The ratio of de novo indels to substitutions, about 1:10, is similar to the ratio we had previously observed over exomes [1].

Although there is no de novo statistical difference between affected and unaffected children for either DIIN or DISB in introns overall, the situation changes if we consider the gene sets enriched in putative ‘autism genes’, the targets of contributory or causal mutation. The statistical significance of delta is very clear for DIIN in the set ‘affected LGD targets’ (Table 1). The delta of 26 events has p-values of .01, .002 and .001 by our three statistical measures. We have estimated that about half of these LGD- target genes are actually autism genes.

In [3], we described a gene protection score that reflects the degree to which disruptive variants in a gene are under strong negative selective pressure in humans. We found evidence that de novo LGDs in protected genes are more likely to be autism genes. We find further evidence for this in the present data. Restricting to the more protected LGD targets, the p-values for the delta gain in significance (p- vals: 0.005, 0.0002, and <0.0001). By contrast, the half of the LGD targets that are less protected show no significant difference as targets for DIIN (p-vals 0.70, 0.24, and 0.37). The delta for the more protected barely shrinks from 26 to 23 while the delta for the less protected shrinks from 26 to 3 (p-val = 0.03 by a permutation test).

In sharp contrast to LGD exon targets in affecteds, we observe no consistent signal for DIIN within gene subsets comprised of de novo LGDs exon targets in siblings, or de novo missense or synonymous substitutions in affected or unaffected siblings. These results are consistent with the hypothesis that there will be little enrichment for autism target genes in these sets. We also observe virtually no signal for DISB for any subset.

### Searching for explanation

None of the events were close to the canonical splice sites: the minimum distance to the site for the de novo indels in affected LGD targets of affected children was 83bp and the majority of events were many kilobases inside the introns (see Table 2). We should note here that the 510 affecteds were chosen to have no mutations of the canonical splice sites that would be observable by exome sequencing. Otherwise we would expect an additional delta of ten de novo events hitting the canonical sites.

**Table 2.**
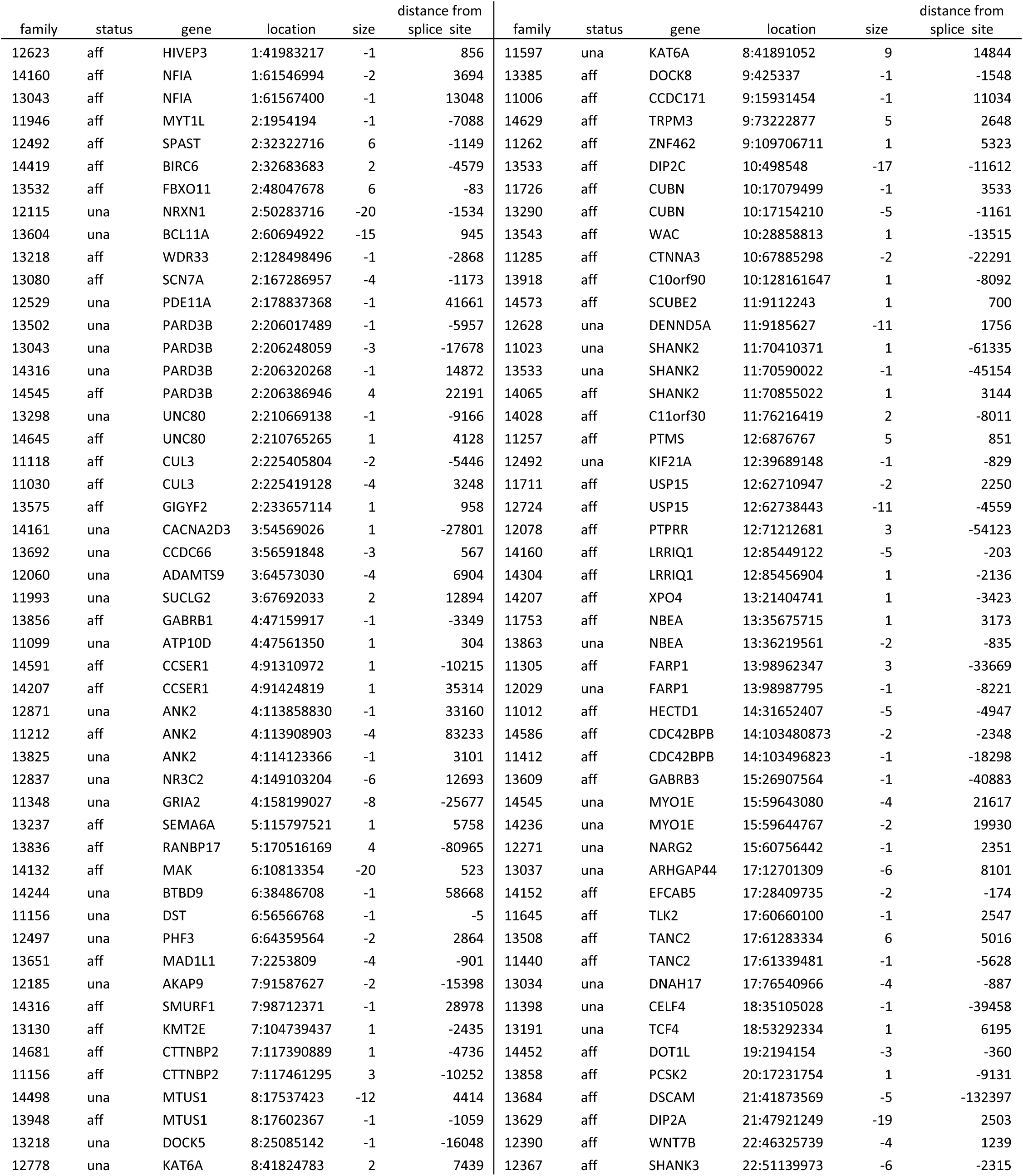
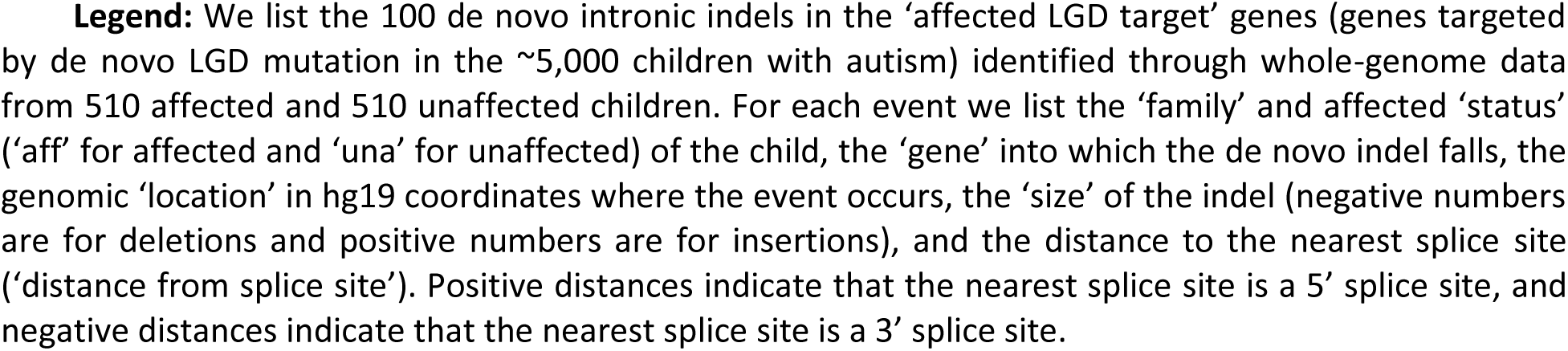
List of de novo intronic indels (DIINs) in the ‘affected LGD targets’

| family | status | gene | location | size | distance from<br>splice site | family | status | gene | location | size | distance from<br>splice site |
| --- | --- | --- | --- | --- | --- | --- | --- | --- | --- | --- | --- |
| 12623 | aff | HIVEP3 | 1:41983217 | -1 | 856 | 11597 | una | KAT6A | 8:41891052 | 9 | 14844 |
| 14160 | aff | NFIA | 1:61546994 | -2 | 3694 | 13385 | aff | DOCK8 | 9:425337 | -1 | -1548 |
| 13043 | aff | NFIA | 1:61567400 | -1 | 13048 | 11006 | aff | CCDC171 | 9:15931454 | -1 | 11034 |
| 11946 | aff | MYT1L | 2:1954194 | -1 | -7088 | 14629 | aff | TRPM3 | 9:73222877 | 5 | 2648 |
| 12492 | aff | SPAST | 2:32322716 | 6 | -1149 | 11262 | aff | ZNF462 | 9:109706711 | 1 | 5323 |
| 14419 | aff | BIRC6 | 2:32683683 | 2 | -4579 | 13533 | aff | DIP2C | 10:498548 | -17 | -11612 |
| 13532 | aff | FBXO11 | 2:48047678 | 6 | -83 | 11726 | aff | CUBN | 10:17079499 | -1 | 3533 |
| 12115 | una | NRXN1 | 2:50283716 | -20 | -1534 | 13290 | aff | CUBN | 10:17154210 | -5 | -1161 |
| 13604 | una | BCL11A | 2:60694922 | -15 | 945 | 13543 | aff | WAC | 10:28858813 | 1 | -13515 |
| 13218 | aff | WDR33 | 2:128498496 | -1 | -2868 | 11285 | aff | CTNNA3 | 10:67885298 | -2 | -22291 |
| 13080 | aff | SCN7A | 2:167286957 | -4 | -1173 | 13918 | aff | C10orf90 | 10:128161647 | 1 | -8092 |
| 12529 | una | PDE11A | 2:178837368 | -1 | 41661 | 14573 | aff | SCUBE2 | 11:9112243 | 1 | 700 |
| 13502 | una | PARD3B | 2:206017489 | -1 | -5957 | 12628 | una | DENND5A | 11:9185627 | -11 | 1756 |
| 13043 | una | PARD3B | 2:206248059 | -3 | -17678 | 11023 | una | SHANK2 | 11:70410371 | 1 | -61335 |
| 14316 | una | PARD3B | 2:206320268 | -1 | 14872 | 13533 | una | SHANK2 | 11:70590022 | -1 | -45154 |
| 14545 | aff | PARD3B | 2:206386946 | 4 | 22191 | 14065 | aff | SHANK2 | 11:70855022 | 1 | 3144 |
| 13298 | una | UNC80 | 2:210669138 | -1 | -9166 | 14028 | aff | C11orf30 | 11:76216419 | 2 | -8011 |
| 14645 | aff | UNC80 | 2:210765265 | 1 | 4128 | 11257 | aff | PTMS | 12:6876767 | 5 | 851 |
| 11118 | aff | CUL3 | 2:225405804 | -2 | -5446 | 12492 | una | KIF21A | 12:39689148 | -1 | -829 |
| 11030 | aff | CUL3 | 2:225419128 | -4 | 3248 | 11711 | aff | USP15 | 12:62710947 | -2 | 2250 |
| 13575 | aff | GIGYF2 | 2:233657114 | 1 | 958 | 12724 | aff | USP15 | 12:62738443 | -11 | -4559 |
| 14161 | una | CACNA2D3 | 3:54569026 | 1 | -27801 | 12078 | aff | PTPRR | 12:71212681 | 3 | -54123 |
| 13692 | una | CCDC66 | 3:56591848 | -3 | 567 | 14160 | aff | LRRIQ1 | 12:85449122 | -5 | -203 |
| 12060 | una | ADAMTS9 | 3:64573030 | -4 | 6904 | 14304 | aff | LRRIQ1 | 12:85456904 | 1 | -2136 |
| 11993 | una | SUCLG2 | 3:67692033 | 2 | 12894 | 14207 | aff | XPO4 | 13:21404741 | 1 | -3423 |
| 13856 | aff | GABRB1 | 4:47159917 | -1 | -3349 | 11753 | aff | NBEA | 13:35675715 | 1 | 3173 |
| 11099 | una | ATP10D | 4:47561350 | 1 | 304 | 13863 | una | NBEA | 13:36219561 | -2 | -835 |
| 14591 | aff | CCSER1 | 4:91310972 | 1 | -10215 | 11305 | aff | FARP1 | 13:98962347 | 3 | -33669 |
| 14207 | aff | CCSER1 | 4:91424819 | 1 | 35314 | 12029 | una | FARP1 | 13:98987795 | -1 | -8221 |
| 12871 | una | ANK2 | 4:113858830 | -1 | 33160 | 11012 | aff | HECTD1 | 14:31652407 | -5 | -4947 |
| 11212 | aff | ANK2 | 4:113908903 | -4 | 83233 | 14586 | aff | CDC42BPB | 14:103480873 | -2 | -2348 |
| 13825 | una | ANK2 | 4:114123366 | -1 | 3101 | 11412 | aff | CDC42BPB | 14:103496823 | -1 | -18298 |
| 12837 | una | NR3C2 | 4:149103204 | -6 | 12693 | 13609 | aff | GABRB3 | 15:26907564 | -1 | -40883 |
| 11348 | una | GRIA2 | 4:158199027 | -8 | -25677 | 14545 | una | MYO1E | 15:59643080 | -4 | 21617 |
| 13237 | aff | SEMA6A | 5:115797521 | 1 | 5758 | 14236 | una | MYO1E | 15:59644767 | -2 | 19930 |
| 13836 | aff | RANBP17 | 5:170516169 | 4 | -80965 | 12271 | una | NARG2 | 15:60756442 | -1 | 2351 |
| 14132 | aff | MAK | 6:10813354 | -20 | 523 | 13037 | una | ARHGAP44 | 17:12701309 | -6 | 8101 |
| 14244 | una | BTBD9 | 6:38486708 | -1 | 58668 | 14152 | aff | EFCAB5 | 17:28409735 | -2 | -174 |
| 11156 | una | DST | 6:56566768 | -1 | -5 | 11645 | aff | TLK2 | 17:60660100 | -1 | 2547 |
| 12497 | una | PHF3 | 6:64359564 | -2 | 2864 | 13508 | aff | TANC2 | 17:61283334 | 6 | 5016 |
| 13651 | aff | MAD1L1 | 7:2253809 | -4 | -901 | 11440 | aff | TANC2 | 17:61339481 | -1 | -5628 |
| 12185 | una | AKAP9 | 7:91587627 | -2 | -15398 | 13034 | una | DNAH17 | 17:76540966 | -4 | -887 |
| 14316 | aff | SMURF1 | 7:98712371 | -1 | 28978 | 11398 | una | CELF4 | 18:35105028 | -1 | -39458 |
| 13130 | aff | KMT2E | 7:104739437 | 1 | -2435 | 13191 | una | TCF4 | 18:53292334 | 1 | 6195 |
| 14681 | aff | CTTNBP2 | 7:117390889 | 1 | -4736 | 14452 | aff | DOT1L | 19:2194154 | -3 | -360 |
| 11156 | aff | CTTNBP2 | 7:117461295 | 3 | -10252 | 13858 | aff | PCSK2 | 20:17231754 | 1 | -9131 |
| 14498 | una | MTUS1 | 8:17537423 | -12 | 4414 | 13684 | aff | DSCAM | 21:41873569 | -5 | -132397 |
| 13948 | aff | MTUS1 | 8:17602367 | -1 | -1059 | 13629 | aff | DIP2A | 21:47921249 | -19 | 2503 |
| 13218 | una | DOCK5 | 8:25085142 | -1 | -16048 | 12390 | aff | WNT7B | 22:46325739 | -4 | 1239 |
| 12778 | una | KAT6A | 8:41824783 | 2 | 7439 | 12367 | aff | SHANK3 | 22:51139973 | -6 | -2315 |

**Legend:** We list the 100 de novo intronic indels in the ‘affected LGD target’ genes (genes targeted by de novo LGD mutation in the ~5,000 children with autism) identified through whole-genome data from 510 affected and 510 unaffected children. For each event we list the ‘family’ and affected ‘status’ (‘aff’ for affected and ‘una’ for unaffected) of the child, the ‘gene’ into which the de novo indel falls, the genomic ‘location’ in hg19 coordinates where the event occurs, the ‘size’ of the indel (negative numbers are for deletions and positive numbers are for insertions), and the distance to the nearest splice site (‘distance from splice site’). Positive distances indicate that the nearest splice site is a 5’ splice site, and negative distances indicate that the nearest splice site is a 3’ splice site.

Almost all the observed indels in affected LGD targets are quite small (see Table 2), with most being of length 1 or 2 nucleotides. The proportion of DIINs with size larger than 2bp in the autism target genes in affected children (25/63 = 40%) is larger than the proportion of such events in the unaffected children (12/37 = 32%) but the difference is not significant by Fisher exact test.

About 10% percent of intronic space falls within 5'UTRs or 3'UTRs. The rest of the introns are between protein coding exons (CDintrons). Significant difference in the delta for DIINs was only seen in the CDintrons, perhaps because of the small size of the former. Table 1 tabulates only de novo events in CDintrons and Supplementary Table 4 tabulates the UTR introns.

In the hope of finding clues to their mechanism of action, we further searched properties of the DIINs. We examined several numerical properties that could reasonably be hypothesized to point to contributory events. These properties were related to the lengths of the affected introns, the proximity of the mutation site to consensus splice sites, the degree of conservation at the mutated site, the likelihood of creation of a new splice site, and the length of the largest open reading frame at that site. The latter might indicate the possibility that the mutation affected an unannotated exon. We associated all de novo intronic events (both indels and substitutions) with each of the above properties, and then asked if the distributions of these properties differed significantly among subsets of the de novo events. These subsets included type (indel or substitution), the affected status of the child, and the target gene class (e.g., ‘all genes’ and ‘affected LGD targets’). None of our efforts were rewarded with a statistically significant signal, but our observations, some positive, are reported in the Supplement.

## Discussion

Once it was shown that germline copy number variation contributes to autism, exome studies became the method of choice to explore germline contribution in greater detail. From exome sequencing, many excellent candidate autism genes have been identified. On the order of 30% of simplex autism is caused in whole or in part by missense, nonsense, splicing or frameshift mutations and large copy number events. Whole genome studies were delayed in part by expense, in part because we cannot predict which noncoding variants alters gene function. However, now that we have good lists of likely autism genes WGS has been performed, in the hopes that statistical signal would emerge by restricting attention to just those genes. There is, moreover, the hope that we will learn which and how noncoding variants alter gene function.

We focused first on intron mutations as there is precedent from previous work that disruption of splicing is frequently a cause for genetic disorders. Although we can infer that the great majority of events within the introns of target genes appear harmless, especially substitutions, we observed a significant excess of de novo indel mutations in affected compared to unaffected siblings. We do not see significant signal for the remainder of the genome, an indication that restricting to likely autism genes matters, and secondarily that the lists of autism genes are good. Autism gene lists further pruned by evidence of negative selective pressure are better still.

Many of the observed de novo indels are only a single nucleotide shift (median = 2, maximum = 47). We see an increase in the indel size in affecteds vs unaffected, but it is not significant. Given the small size of indels, we were a little surprised to see no significant signal coming from de novo substitution events in those introns. However, de novo substitutions are ten times more common than indels, and a larger proportion of substitutions are likely to be harmless, so signal from them is more likely to be hidden in noise. Additionally, an indel could potentially cause a substantial alteration in the conformation of RNA or DNA that may propagate for several nucleotides, or perhaps longer, creating a structure that might not be recognized by a binding protein, whereas the effect of a substitution is more likely to be very local.

Our entire signal falls within the introns between coding exons. We infer from this that they do indeed disrupt splicing, but we have no direct demonstration of this. All of our attempts to find statistical evidence for known molecular mechanisms yielded nothing of significance. The indels are generally deep within the introns. Not only do they not occur at the consensus splice sites, but they are far clear of them. They do not appear to create new 3’ or 5’ splice sites, nor disrupt cryptic open reading frames, nor disrupt any of the highly conserved elements within introns identified through comparative genomics. So, although the introns appear to be full of sensitive “targets”, we fail to see a predominant explanation, one that yields statistical significance. We feel that how these mutations act is now an open question. Are they interfering with splicing, or targeting control regions? This uncertainty invites future attention as we try to understand the molecular biology of the gene.

We are also now in a position to better estimate the overall contribution of germline mutation to autism diagnosis. 26 more intronic indels occur within the 546 LGD target genes (Table 1) in the affected vs unaffected. There are 510 discordant siblings, so we infer that as many as 5% (26/510) have a diagnosis of autism in part due to de novo intronic indels. From the whole-exome studies we have estimated that only about half of the affected LGD targets are true autism genes and that the number of true autism genes is about 500. These enable us to extrapolate as many as ∼10% of the SSC children would have autism due to de novo intronic indels in autism genes. The observed delta of 61 of de novo intronic events in all genes supports that extrapolation. It is almost assured that other de novo intronic events like substitutions, microsatellite expansions, and indels of sizes larger than we can presently detect also contribute to the disorder. If such presently cryptic events contributed in an amount about equal to small de novo indels in introns, the total contribution would be about ∼20%. This figure is only slightly less than our estimates of the contribution from de novo missense, nonsense, and frame-shifts combined. If indeed most harmful intron mutations disturb splicing, altered splicing is a very major cause of genetic abnormalities.

Assuming contributions of de novo coding mutations (∼20%), de novo intronic events (∼20%) and de novo CNV (∼6%) the combination is about 46%, bringing us very close to our theoretical expectation of 60% contribution for de novo germline mutations in simplex autism [17]. The remaining gap might be filled by de novo mutation in intergenic control regions or in noncoding transcripts or in the long range effects of rearrangements that we do not yet identify.

## Acknowledgments

We thank all the families at the participating SSC sites, as well as the principal investigators (A. L. Beaudet, R. Bernier, J. Constantino, E. H. Cook, Jr., E. Fombonne, D. Geschwind, D. E. Grice, A. Klin, D. H. Ledbetter, C. Lord, C. L. Martin, D. M. Martin, R. Maxim, J. Miles, O. Ousley, B. Peterson, J. Piggot, C. Saulnier, M. W. State, W. Stone, J. S. Sutcliffe, C. A. Walsh, and E. Wijsman) and the coordinators and staff at the SSC sites for the recruitment and comprehensive assessment of simplex families, and the Simons Foundation Autism Research Initiative (SFARI) staff for facilitating access to the SSC. This work was supported by SFARI Grants SF235988 (to M.W.) and SF362665 (to I.I.) and NIH Grant NIH 1UM1HG008901-01 (to RB.D.).

## Supplement Methods

### Measuring significance of delta

There are three different methods for testing if the number of de novo intronic events in affected and unaffected children is significantly different than the expectation of equality.

#### Chi square test

This test compares the two de novo intronic event numbers in affected vs. unaffected children for a given target gene class (e.g., ‘affected LGD targets’) to the two event numbers for ‘all genes’ in DISB.

#### Status permutation method

It is a permutation test based on random swapping of the number of de novo intronic events for the discordant sibling pairs (affected vs. unaffected) for a given target gene class.

#### Gene permutation method

It measures the significance of observed difference in the number of de novo intronic events in affected and in unaffected children. In this method, we select genes with similar intron lengths as the genes in the analyzed gene set. As a measure of intronic lengths we used the number of ultra-rare substitutions (variants seen only once in the 1020 parents). The total length of the introns in a gene (measured using RefSeq gene model databases) and the number of ultra-rare intronic substitutions are linearly related, but we chose to use the number of intronic substitutions because it accounts for the coverage in the whole-genome data (Table S3 shows the intron lengths and the number of ultra-rare substitutions for each gene).

To select random gene set of genes with similar number of ultra-rare intronic substitutions as the analyzed set, we first sorted all the genes based on the number of ultra-rare intronic substitutions. Then for each of the analyzed genes we selected randomly either the previous or the following gene from the sorted list of genes.

### Searching for explanation

We observed that in the affected children there were significantly more de novo intronic indels in the autism targets genes than in the unaffected children. We inferred that the increase is due to the indirect ascertainment of intronic indels that contributed to diagnosis of autism in the affected children and we asked the natural question if the contributory de novo intronic indels could be distinguished from the non-contributory events by some of their properties. We examined 15 numerical properties (see the detailed list and description below) that could reasonably be hypothesized to point to contributory events. We associated all de novo intronic events (both indels and substitutions) with each of the 15 properties and tested if the distributions of these properties differed among subsets of the de novo events defined by the de novo intronic event type (indel or substitution), the affected status of the child carrying the de novo events (affected or unaffected) and by the class of the gene targeted by the event (‘all genes’ or ‘autism target genes’). We performed three different comparisons over the distributions of each property for the subsets of de novo intronic indels: the distribution for all de novo intronic events in affected children vs the distribution for all de novo intronic events in unaffected children (designated as ‘(all, aff) vs (all, una)’); the distribution of the de novo intronic events in the affected children that fall in the autism target genes vs the distribution of all de novo intronic events in the affected children (‘(tar,aff) vs (all,aff)’); the distribution of de novo intronic indels in the target genes in affected children vs the events in target genes in unaffected children (‘(tar,aff) vs (tar,una)’). We also performed the corresponding tests for the de novo intronic substitutions and the six p-values computed using ranksum tests for all properties are shown in Table S5. More detailed view of the distributions of each of the properties over the various classes of events can be seen in the Supplementary Figures 3-17.

### Properties

#### Intron length and distance to the nearest splice-site

For every de novo intronic variant we identified the shortest intron covering the variant. We recorded the length of the shortest intron (‘intron length’ property; see Table S4). We also recorded the distance between the de novo event and the splice-sites of the shortest intron that was closest to the observed event (‘distance from splice-site’ property). We assigned positive number if the closer splice-site was the donor splice-site and negative number if the closer splice-site was the acceptor splice-site. We tested if the absolute value of the distance from splice-site was different between the various classes of the de novo mutations (Figure S3).

#### Open Reading Frame length

To test if the de novo intronic events fall in and disrupted cryptic coding exons, we looked for a bias in the size of the largest open reading frame in the direction of transcription (see ‘ORF length’ property’) among the difference lasses of de novo events (Figure S5).

#### Conservation scores

We used two methods for measuring conservation: phastCons [1] and phyloP [2]. The two methods compute a conservation score for each genomic location based on a given phylogenetic three. We downloaded the computed scores from the two methods over three different phylogenetic trees: vertebrate, placental, and primates from UCSC genome browser. (Figures S12-S17).

#### Novel splice site scores

To test if the de novo intronic mutations created novel splice sites we developed a donor and an acceptor splice-site sequence scores for a given short sequence (see below for detailed definition of the scores). We computed these two scores for the reference sequence around (5 bases up and downstream) the location where the de novo event occurred (‘ref’ scores) and separately for the local sequence after the de novo event was introduced (‘alt’ scores). We also computed the differences between the ‘alt’ scores and the ‘ref’. Thus, every de novo intronic mutation was associated with six splice-site sequence scores: ‘ref’, ‘alt’, ‘alt-ref’ for both donor and acceptor splice-site scores (Tables S2 and S3) and we tested each of the six scores for their ability to separate de novo intronic events in affected children in target genes (Supplementary Table 5 and Supplementary Figures 6-11).

### Definition of the donor and acceptor splice-site sequence scores

We defined a position-specific sequence models for donor and acceptor splice sites based on 20bp sequence context (10bp upstream and 10bp downstream of the splice site). We measured the frequency of the four nucleotides at each of the 20 positions independently using the ∼200,000 annotated donor 𝒜 and acceptor sites in the RefSeq database: 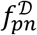 and 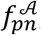, where 𝒟 is for donor, 𝒜 is for acceptor, p is index for the position and n is A, C, G, or T. We also measured the frequency of the random intronic ℛ nucleotides, 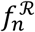 ratios: and defined the position specific donor and acceptor splice-site scores as log-likelihood

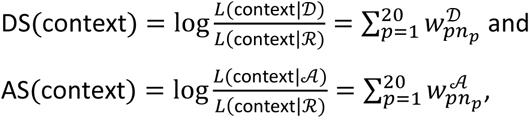

where ‘context’ is the 20bp sequence context around a candidate splice-site position, L(context|M) is the likelihood function for the context given a specified model M under the assumption of independence among the context positions, *n_p_* is the p-th nucleotide in context, 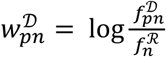, and 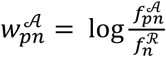

(Supplementary Figure 1).

Finally, we defined the donor and acceptor splice-site sequence scores for a given short sequence, seq, as the maximum of the position-specific splice-site scores over all positions in seq:

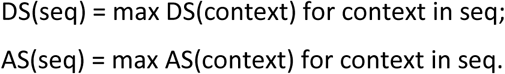

See Supplementary Figure 2 for example AS score for the ‘ref’ and ‘alt’ score for a de novo intronic insertion.

## Supplementary Tables

**Table S1 and S2: Lists of de novo intronic indels (S1) and substitutions (S2)**

The two tables S1 (Supp-T1-DN-indel.xlsx data file) and S2 (Supp-T2-DN-sub.xlsx data file) list all analyzed de novo intronic events, 2,231 indels and 23,715 substitutions, respectively. For each event the tables lists: the ‘family’ and the child (‘in child) where the de novo events are found (prb – is the proband or affected child, sib is for the unaffected sibling, M for male and F for female; some events are shared between the two siblings); the detail description of the variant using VCF conventions (‘variant’ with <chr>:<pos>:<reference allele>:<alternative allele> format, the location <chr>:<pos> in hg19 coordinates) and the ‘variant size’ (0 for substitutions, negative number for deletion and positive number for insertions); the ‘gene’ affected by the variant and the ‘variant effect’ (CDintron for coding introns, 5Uintrons or 3Uintrons). The table also shows if the affected gene is a member of one of the 8 analyzed gene classes (the purple columns) and the 15 analyzed properties of de novo intronic events (blue columns). See Supplementary methods for a description of those properties.

**Table S3: Gene Table**

This table is in the Supp-T3-genes.xlsx data file and shows information about the 23,953 annotated human genes. For each gene, the table lists the ‘gene’ name, gene protection information as reported in [3] (red columns); lengths of the intronic space for each of the three classes of introns computed from the RefSeq gene model database (blue columns); the number of ultra-rare (UR) events by type of the events (sub for substitution, del for deletion, ins for insertion) and by the type of the affected intron (CDintron, 5Uintron, or 3Uintron) (yellow columns); the number of de novo intronic events by the affected status of the child, the type of de novo event and by the type of the affected intron (green columns); and the membership of the gene in each of the 8 genes sub-classes defined by the affected ‘status’ of the child carrying the de novo events (affected or unaffected), by the effect of the de novo event, and based on the degree of protection of the affected gene (purple columns).

**Table S4.**
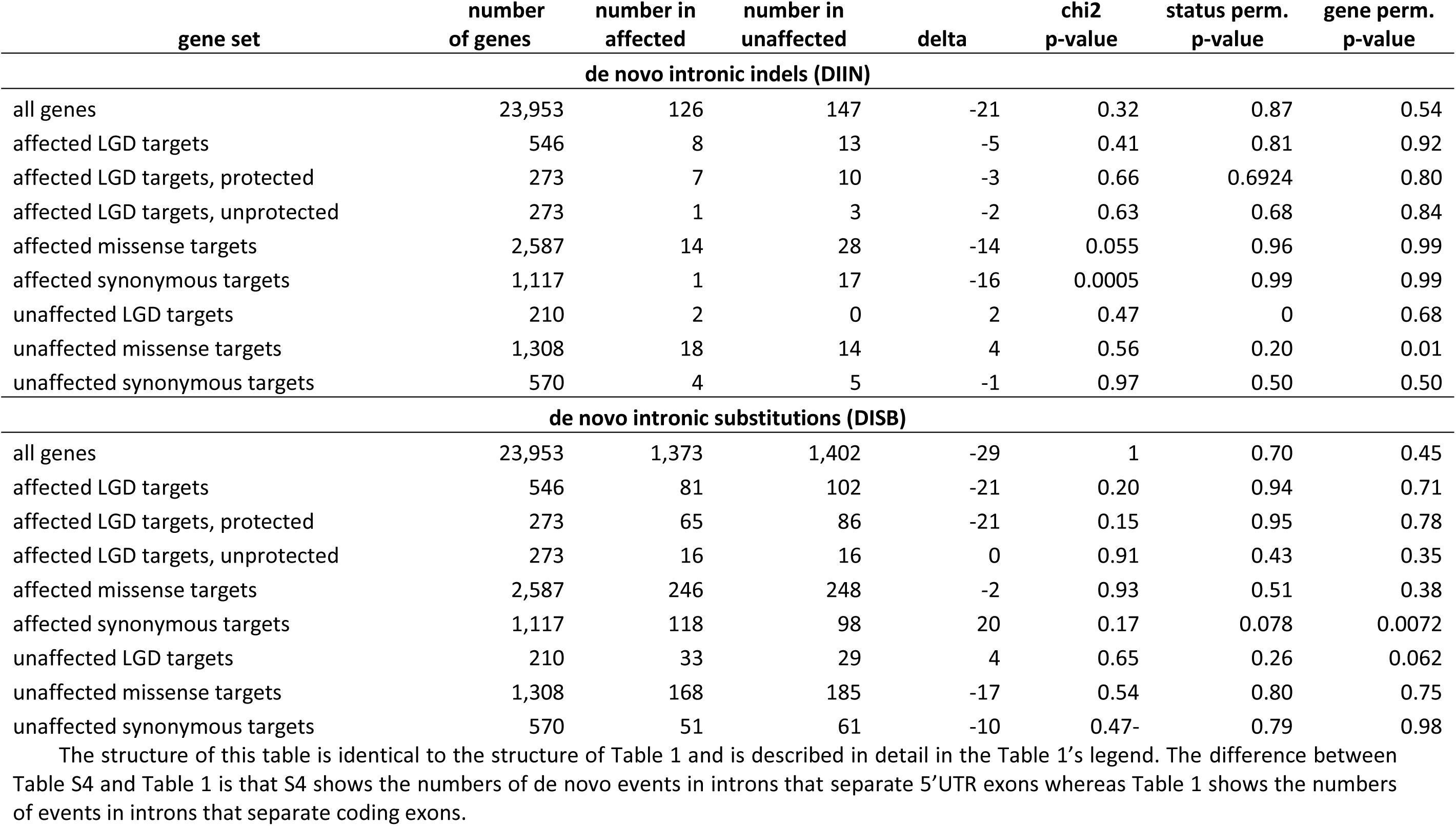
De novo intronic indels (DIIN) and substitutions (DISB) in introns between 5'UTR exons

**Table S5.**
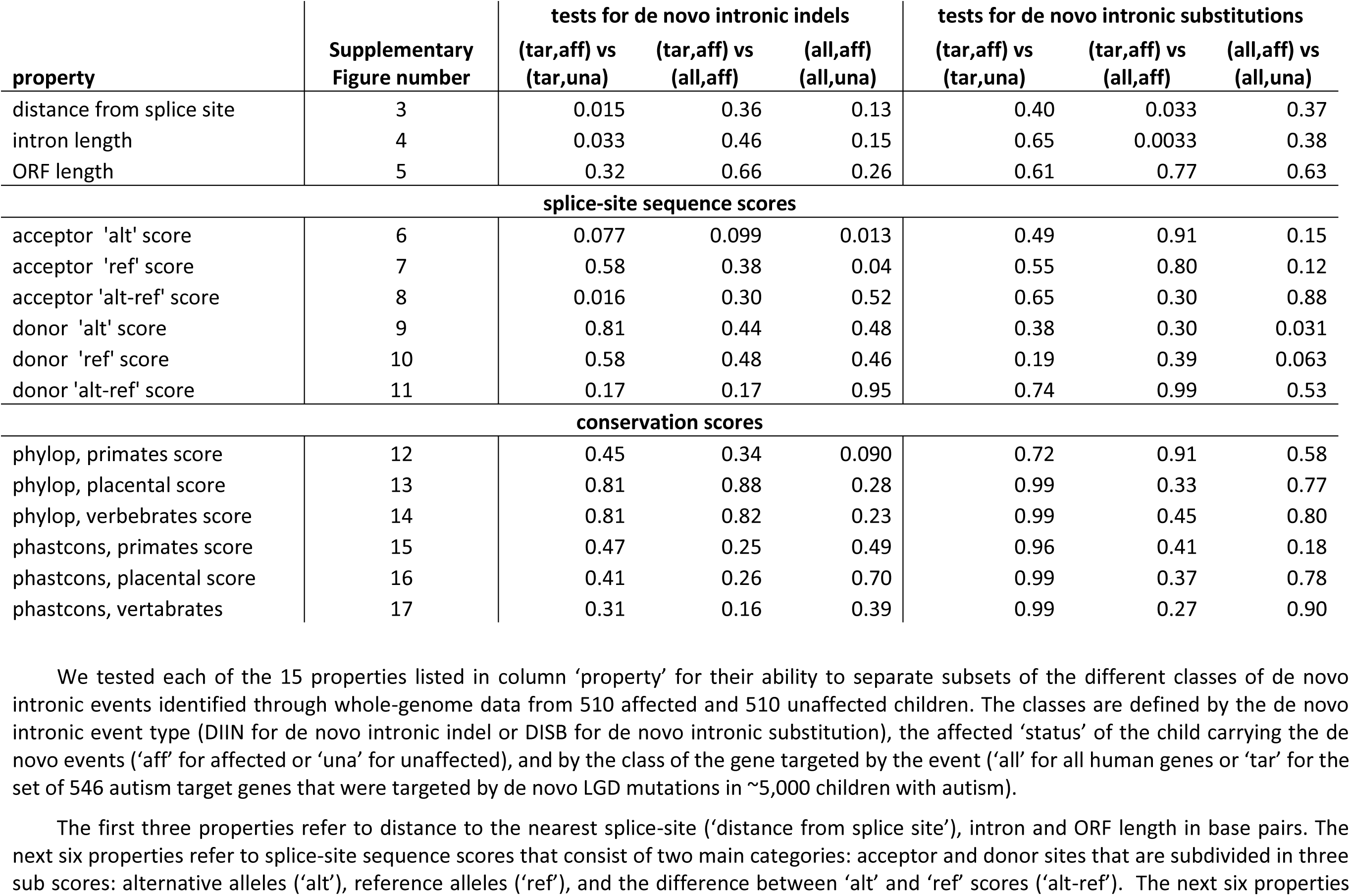
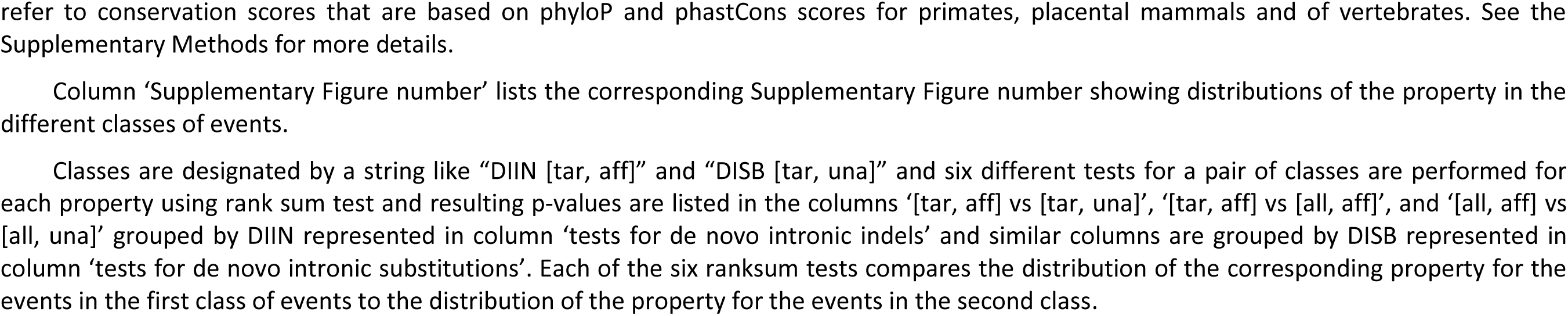
Property Table

| property | Supplementary<br>Figure number | tests for de novo intronic indels |  |  | tests for de novo intronic substitutions |  |  |
| --- | --- | --- | --- | --- | --- | --- | --- |
|  |  | (tar,aff) vs<br>(tar,una) | (tar,aff) vs<br>(all,aff) | (all,aff)<br>(all,una) | (tar,aff) vs<br>(tar,una) | (tar,aff) vs<br>(all,aff) | (all,aff) vs<br>(all,una) |
| distance from splice site | 3 | 0.015 | 0.36 | 0.13 | 0.40 | 0.033 | 0.37 |
| intron length | 4 | 0.033 | 0.46 | 0.15 | 0.65 | 0.0033 | 0.38 |
| ORF length | 5 | 0.32 | 0.66 | 0.26 | 0.61 | 0.77 | 0.63 |
| **splice-site sequence scores** |  |  |  |  |  |  |  |
| acceptor 'alt' score | 6 | 0.077 | 0.099 | 0.013 | 0.49 | 0.91 | 0.15 |
| acceptor 'ref' score | 7 | 0.58 | 0.38 | 0.04 | 0.55 | 0.80 | 0.12 |
| acceptor 'alt-ref' score | 8 | 0.016 | 0.30 | 0.52 | 0.65 | 0.30 | 0.88 |
| donor 'alt' score | 9 | 0.81 | 0.44 | 0.48 | 0.38 | 0.30 | 0.031 |
| donor 'ref' score | 10 | 0.58 | 0.48 | 0.46 | 0.19 | 0.39 | 0.063 |
| donor 'alt-ref' score | 11 | 0.17 | 0.17 | 0.95 | 0.74 | 0.99 | 0.53 |
| **conservation scores** |  |  |  |  |  |  |  |
| phylop, primates score | 12 | 0.45 | 0.34 | 0.090 | 0.72 | 0.91 | 0.58 |
| phylop, placental score | 13 | 0.81 | 0.88 | 0.28 | 0.99 | 0.33 | 0.77 |
| phylop, vertebrates score | 14 | 0.81 | 0.82 | 0.23 | 0.99 | 0.45 | 0.80 |
| phastcons, primates score | 15 | 0.47 | 0.25 | 0.49 | 0.96 | 0.41 | 0.18 |
| phastcons, placental score | 16 | 0.41 | 0.26 | 0.70 | 0.99 | 0.37 | 0.78 |
| phastcons, vertebrates | 17 | 0.31 | 0.16 | 0.39 | 0.99 | 0.27 | 0.90 |

We tested each of the 15 properties listed in column 'property' for their ability to separate subsets of the different classes of de novo intronic events identified through whole-genome data from 510 affected and 510 unaffected children. The classes are defined by the de novo intronic event type (DIIN for de novo intronic indel or DISB for de novo intronic substitution), the affected 'status' of the child carrying the de novo events ('aff' for affected or 'una' for unaffected), and by the class of the gene targeted by the event ('all' for all human genes or 'tar' for the set of 546 autism target genes that were targeted by de novo LGD mutations in ~5,000 children with autism).
The first three properties refer to distance to the nearest splice-site ('distance from splice site'), intron and ORF length in base pairs. The next six properties refer to splice-site sequence scores that consist of two main categories: acceptor and donor sites that are subdivided in three sub scores: alternative alleles ('alt'), reference alleles ('ref'), and the difference between 'alt' and 'ref' scores ('alt-ref'). The next six properties
refer to conservation scores that are based on phyloP and phastCons scores for primates, placental mammals and of vertebrates. See the Supplementary Methods for more details.
Column 'Supplementary Figure number' lists the corresponding Supplementary Figure number showing distributions of the property in the different classes of events.
Classes are designated by a string like "DIIN [tar, aff]" and "DISB [tar, una]" and six different tests for a pair of classes are performed for each property using rank sum test and resulting p-values are listed in the columns '[tar, aff] vs [tar, una]', '[tar, aff] vs [all, aff]', and '[all, aff] vs [all, una]' grouped by DIIN represented in column 'tests for de novo intronic indels' and similar columns are grouped by DISB represented in column 'tests for de novo intronic substitutions'. Each of the six ranksum tests compares the distribution of the corresponding property for the events in the first class of events to the distribution of the property for the events in the second class.

## Supplementary Figures

**Figure S1:**
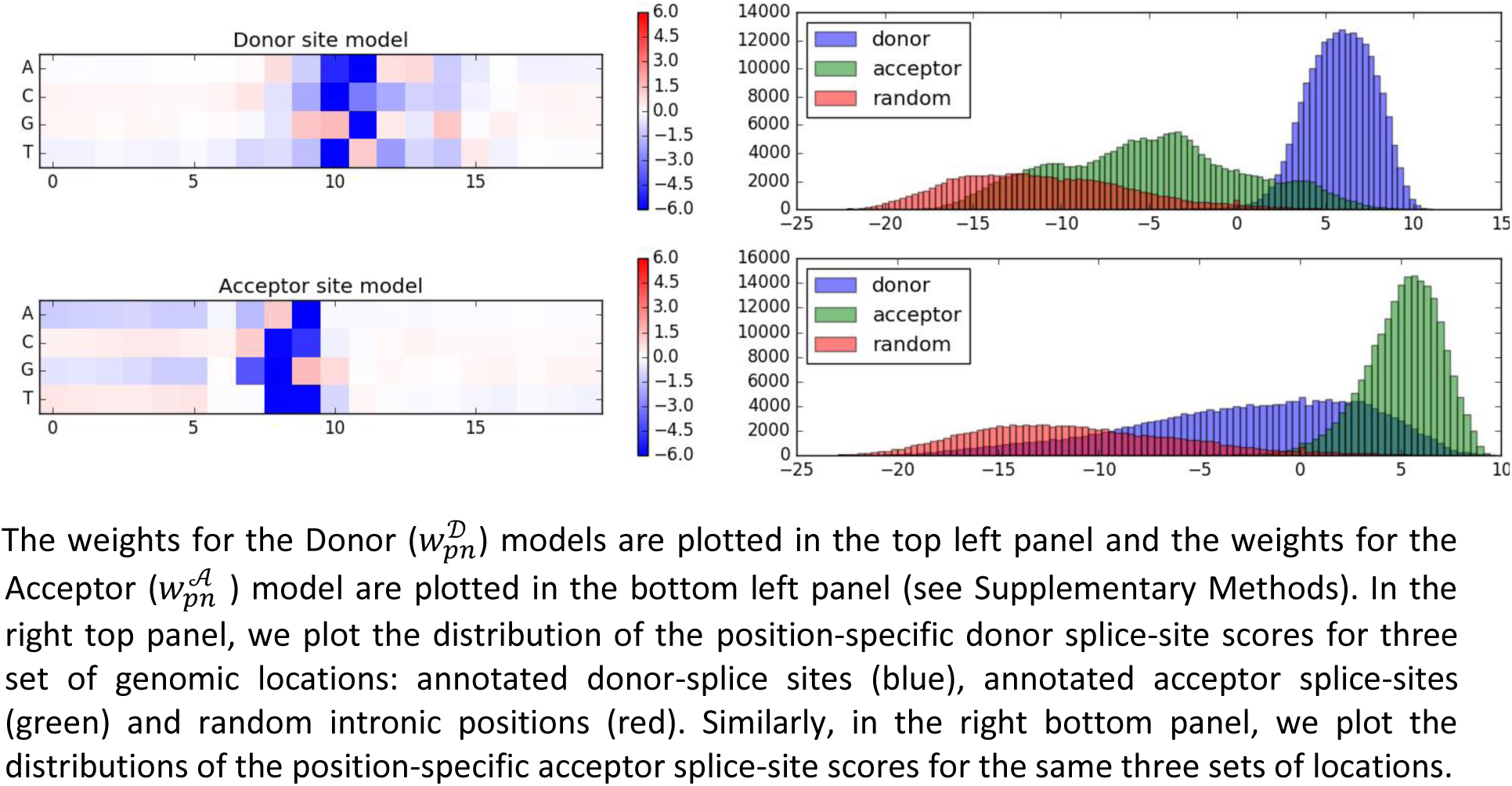
Donor and Acceptor splice-site models

**Figure S2:**
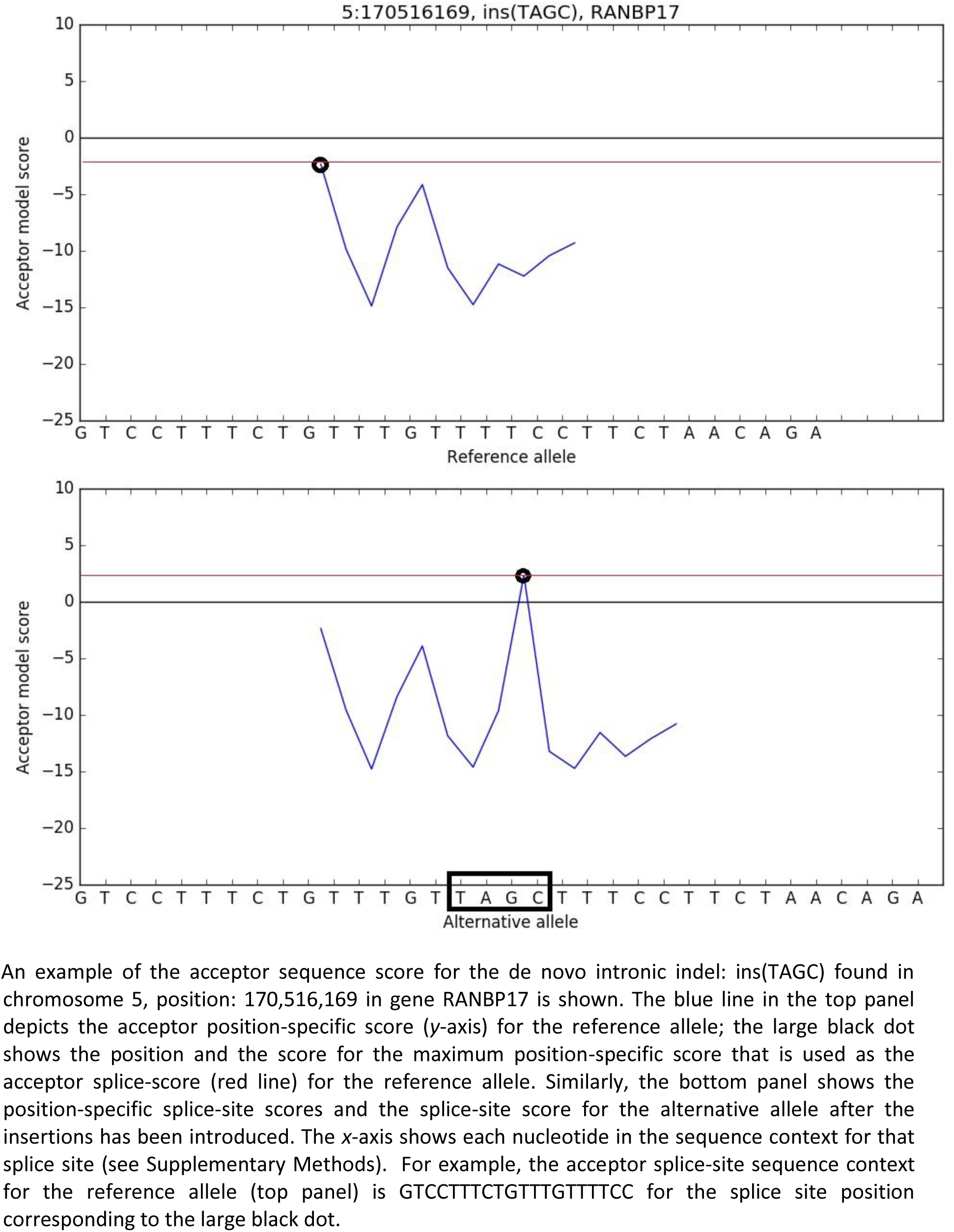
An example of acceptor splice-site sequence score

**Figure S3.**
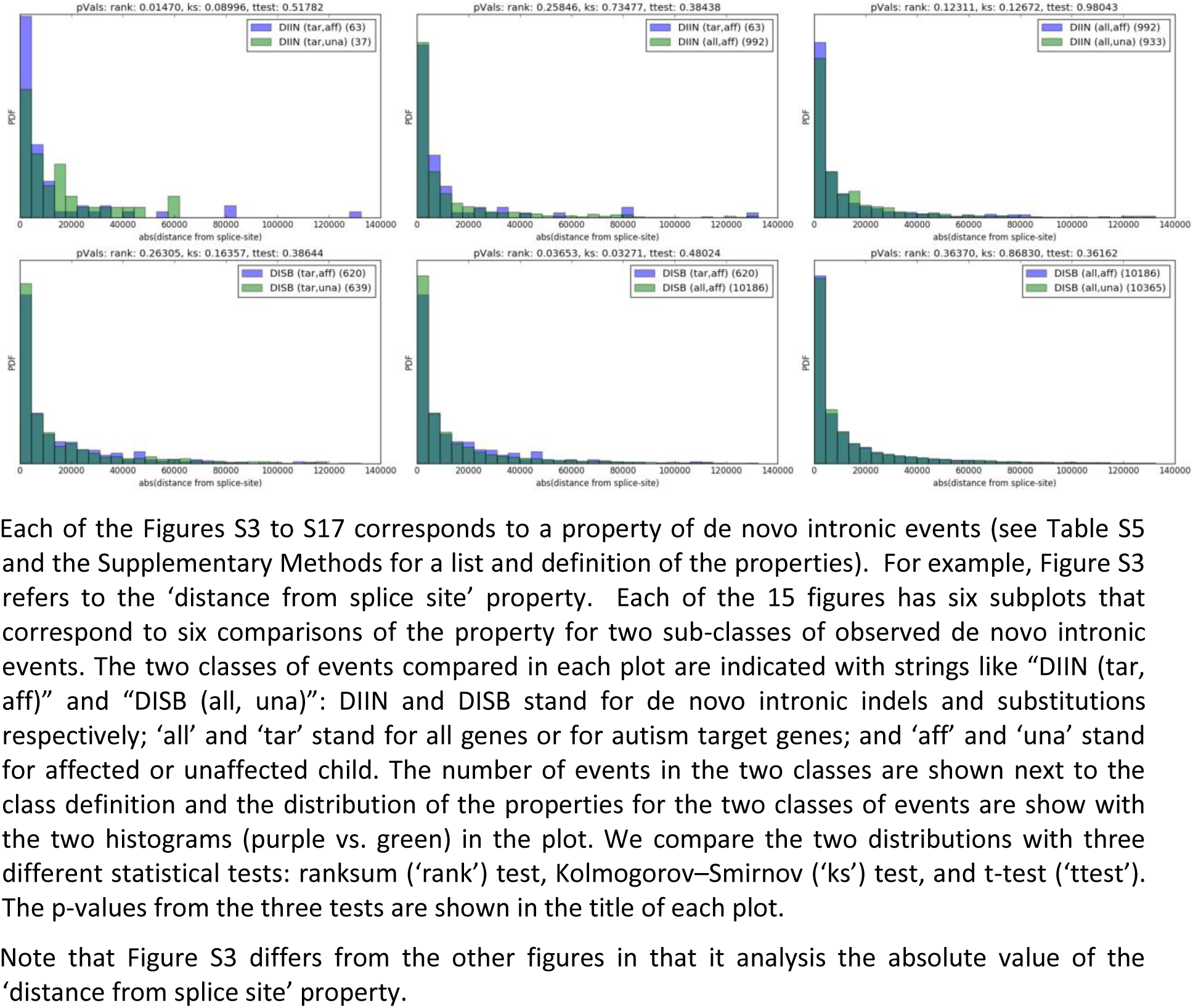
Distance from splice site distributions

**Figure S4.**
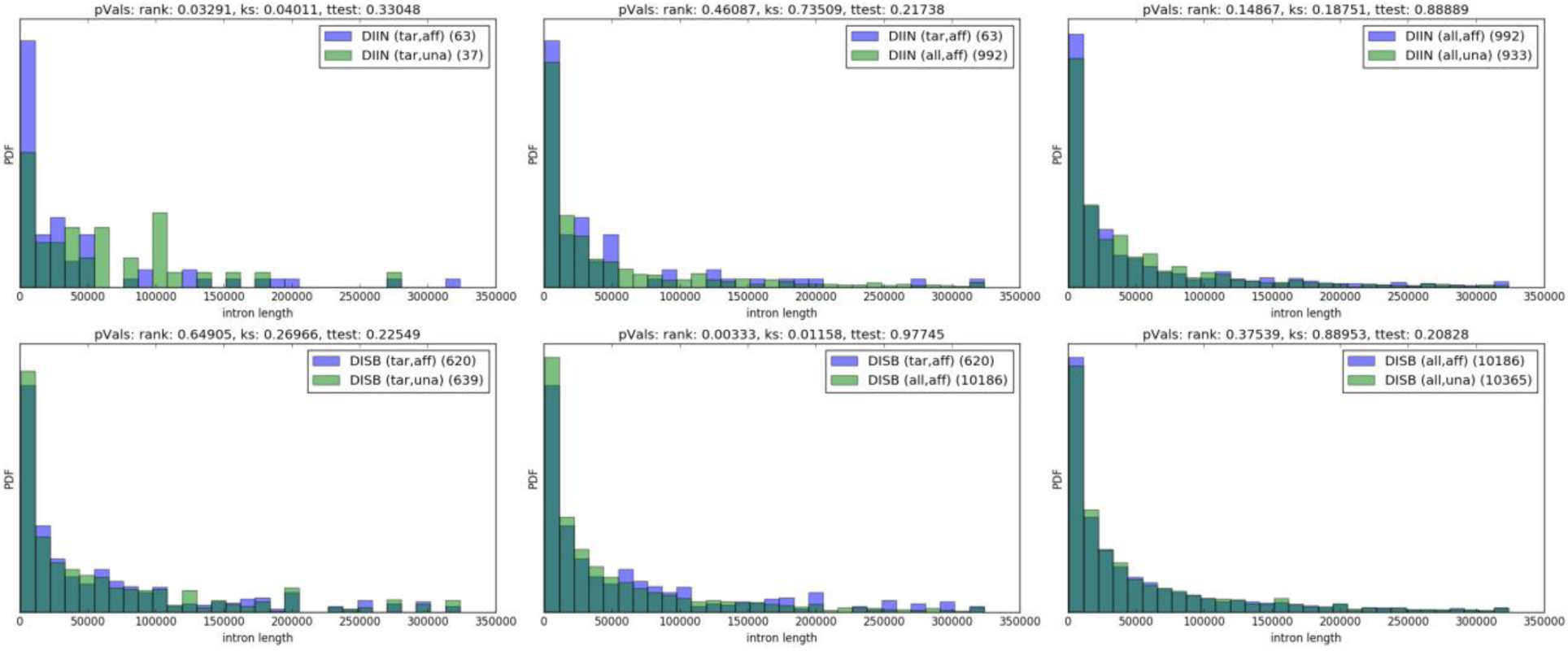
intron length distributions

**Figure S5.**
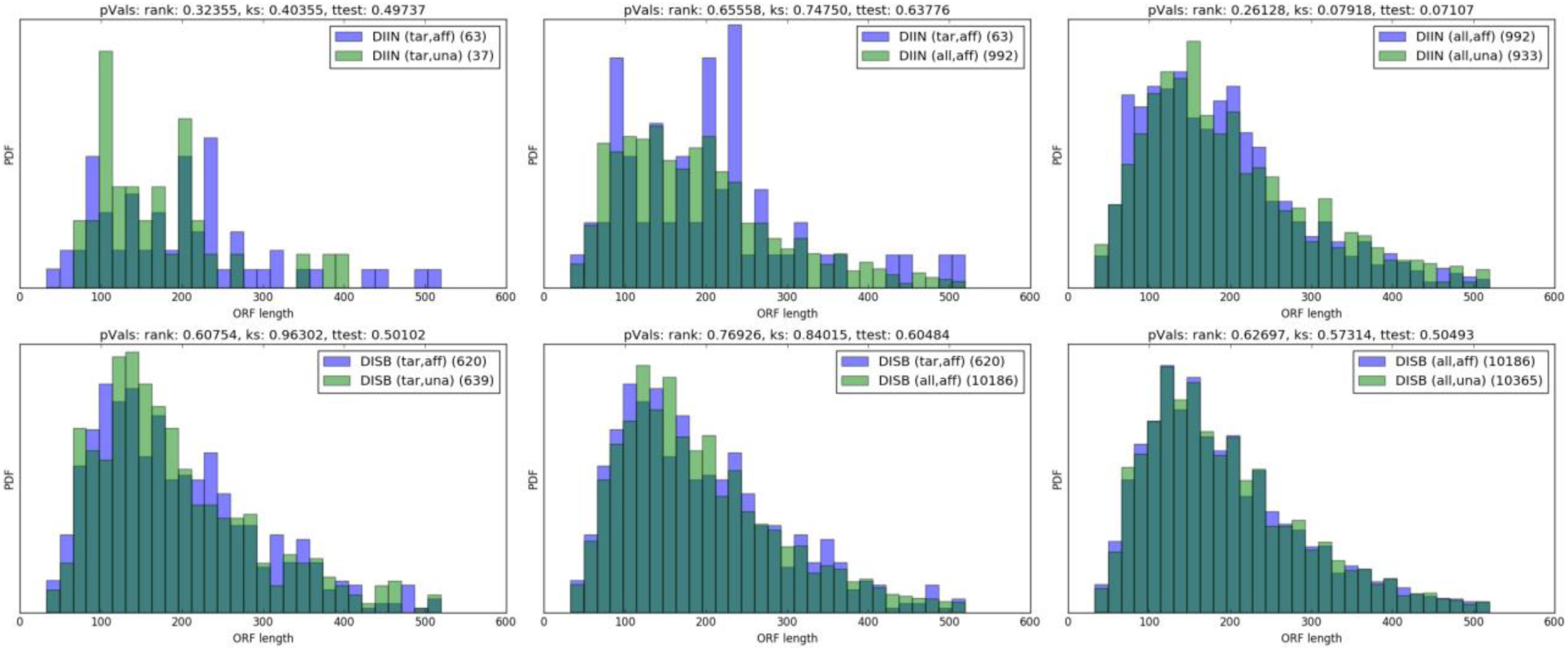
ORF length distributions

**Figure S6.**
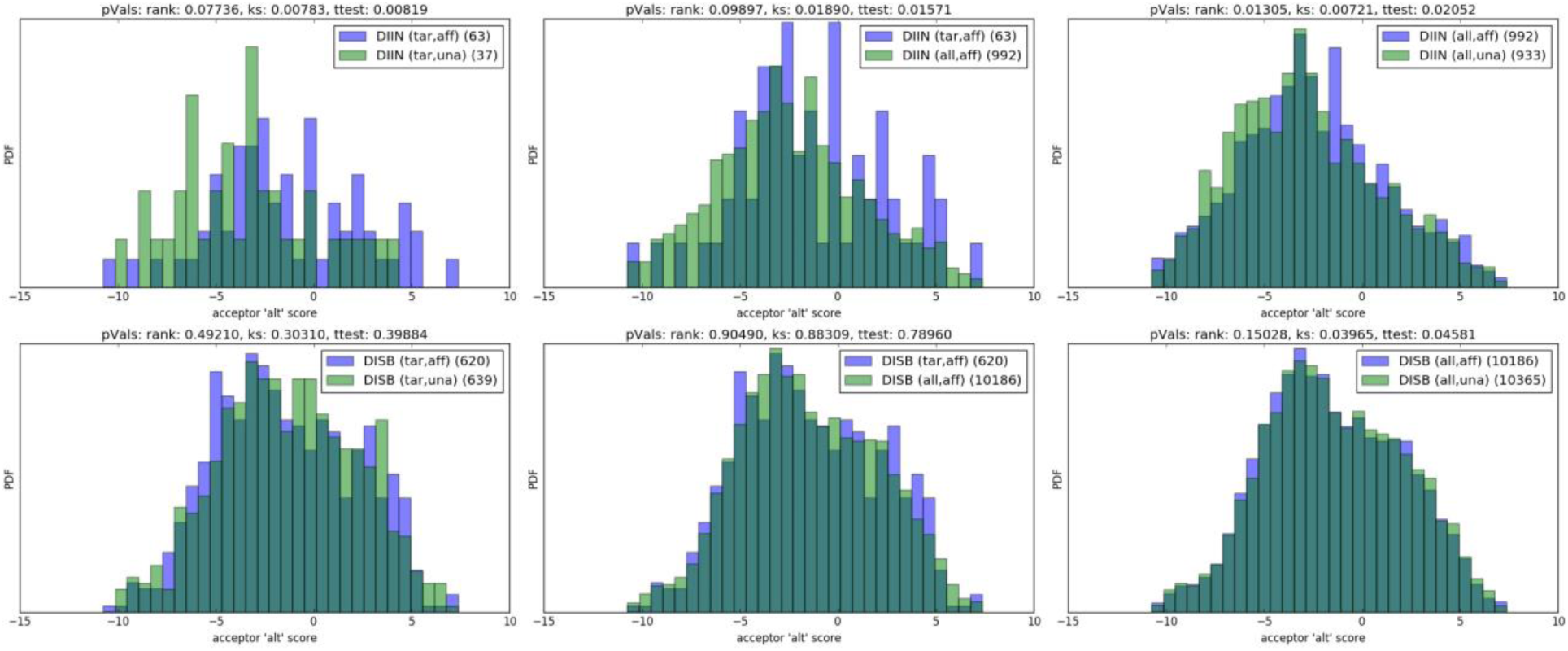
acceptor ‘alt‘ score distributions

**Figure S7.**
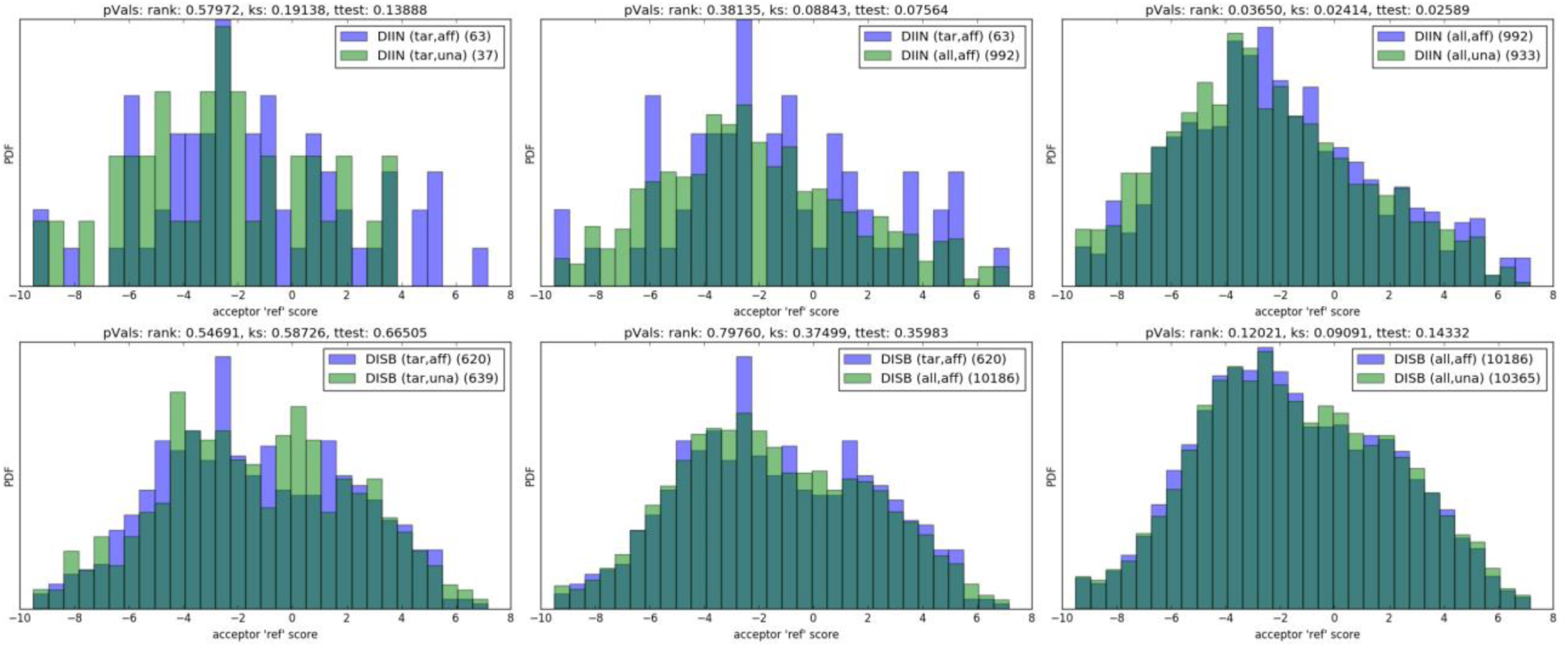
acceptor 'ref' score distribuions

**Figure S8.**
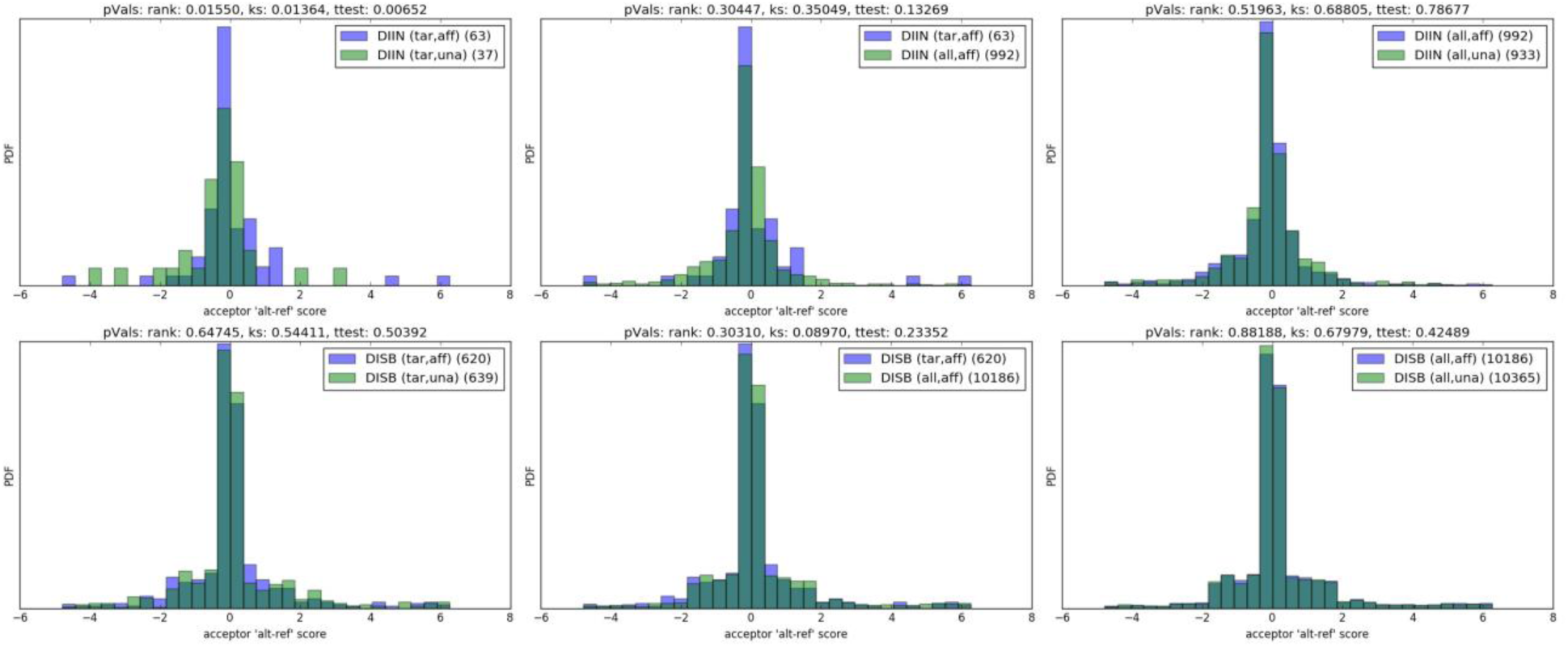
acceptor ‘alt-ref’ score distributions

**Figure S9.**
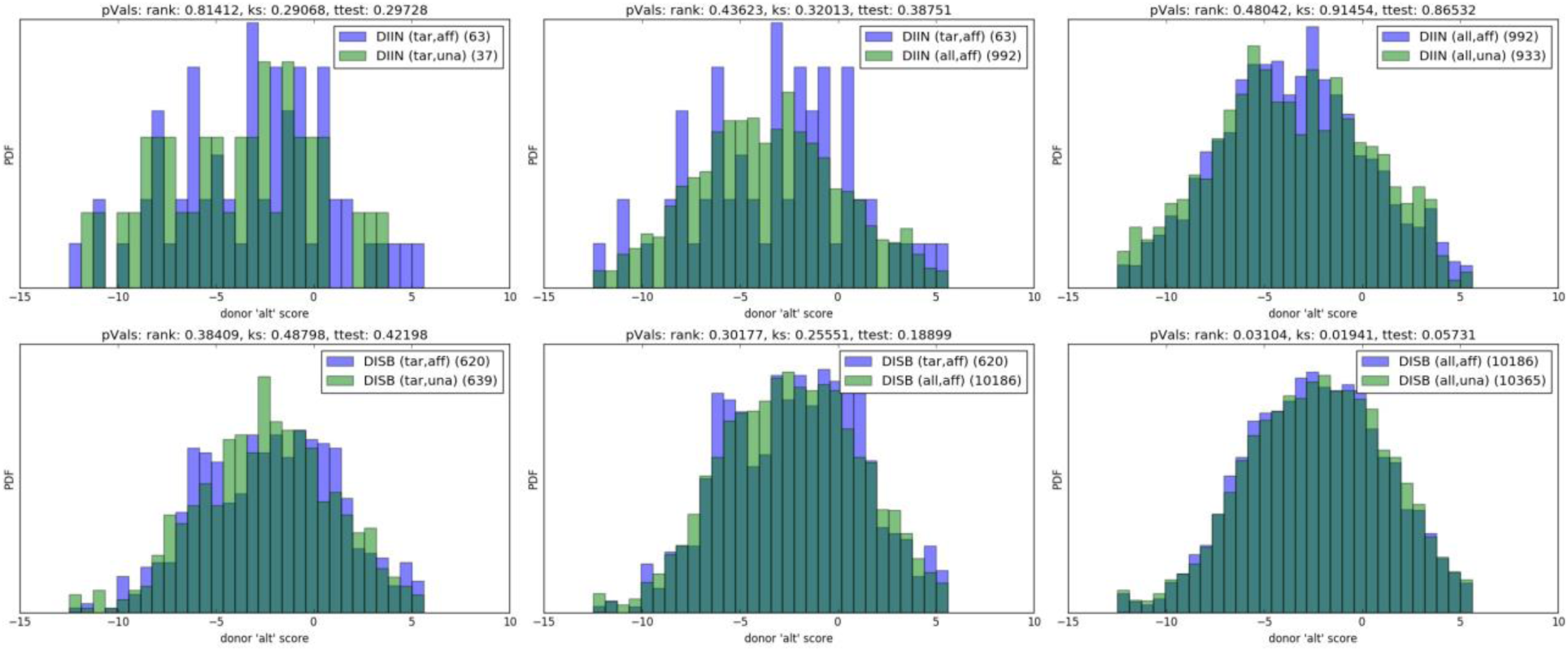
donor 'alt' score distributions

**Figure S10.**
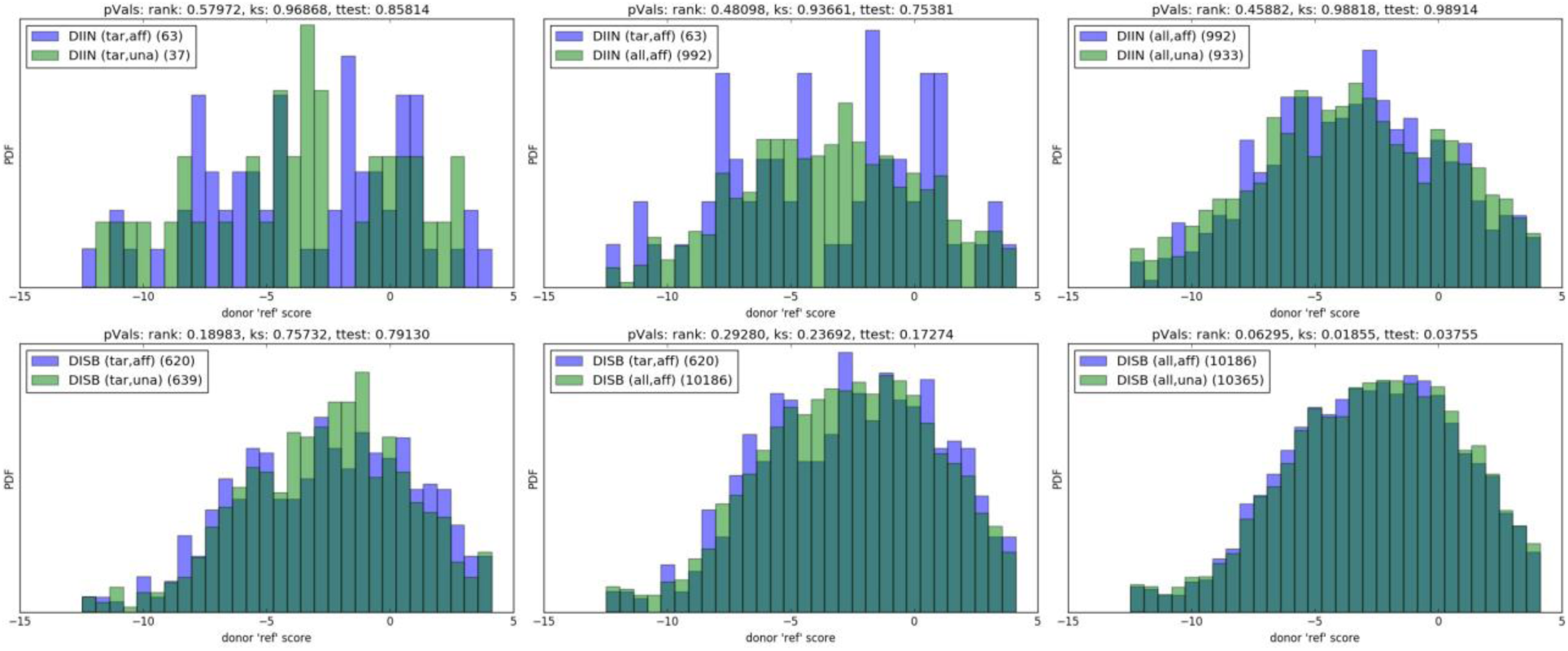
donor ‘ref‘ score distributions

**Figure S11.**
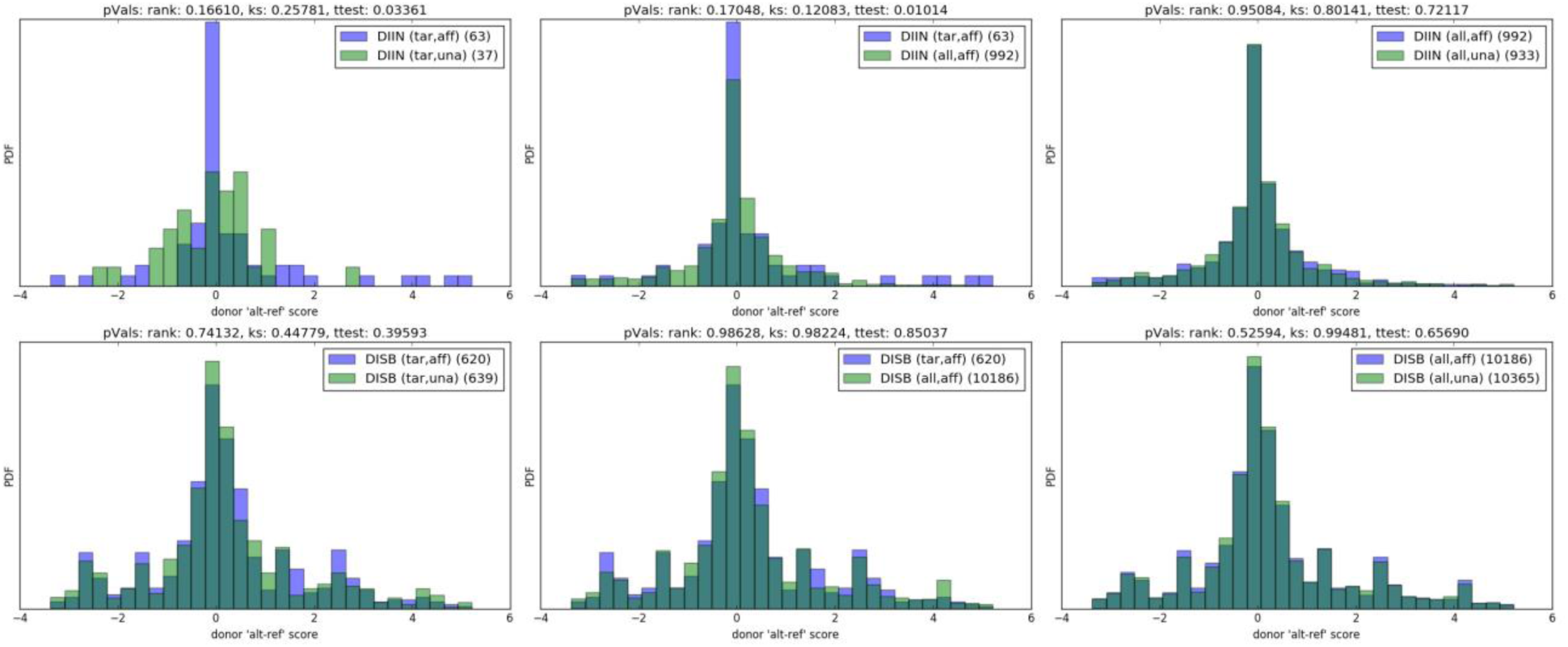
donor 'alt-ref' score distributions

**Figure S12.**
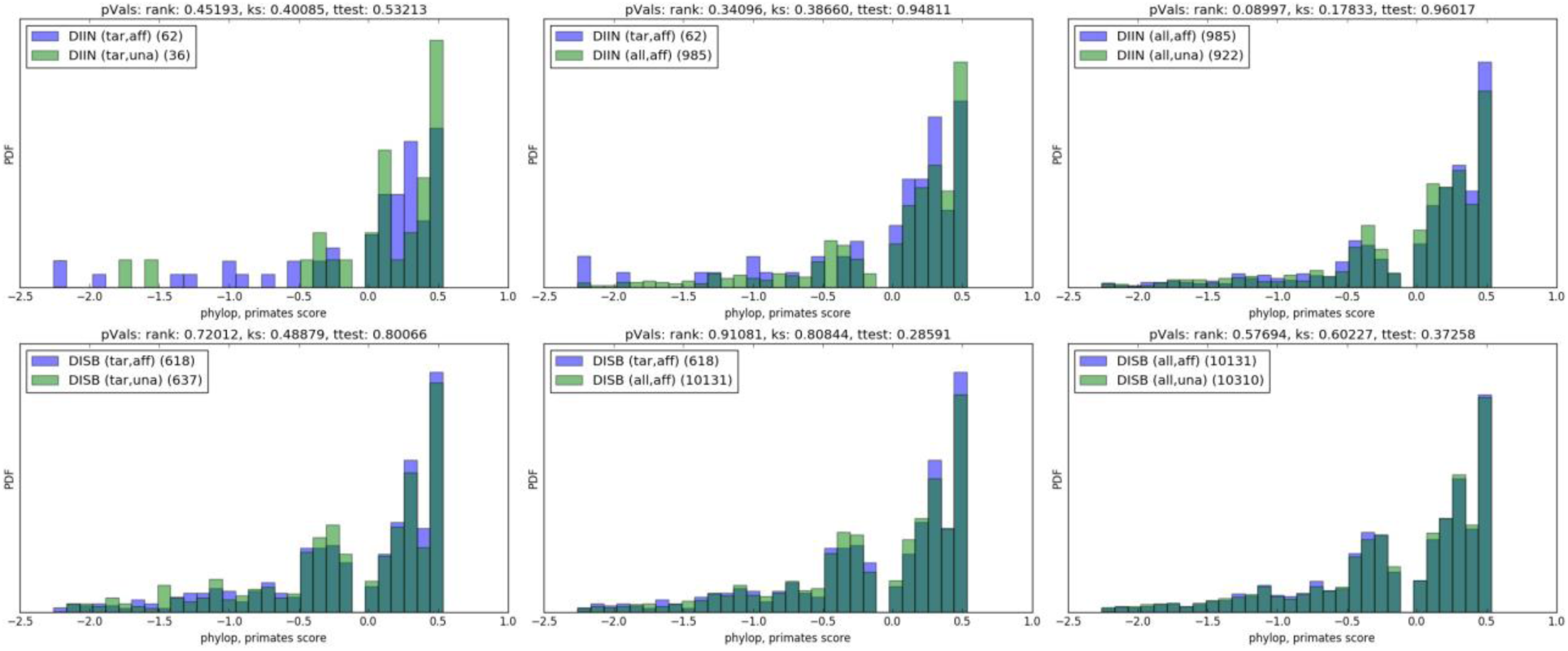
phylop, primates score distributions

**Figure S13.**
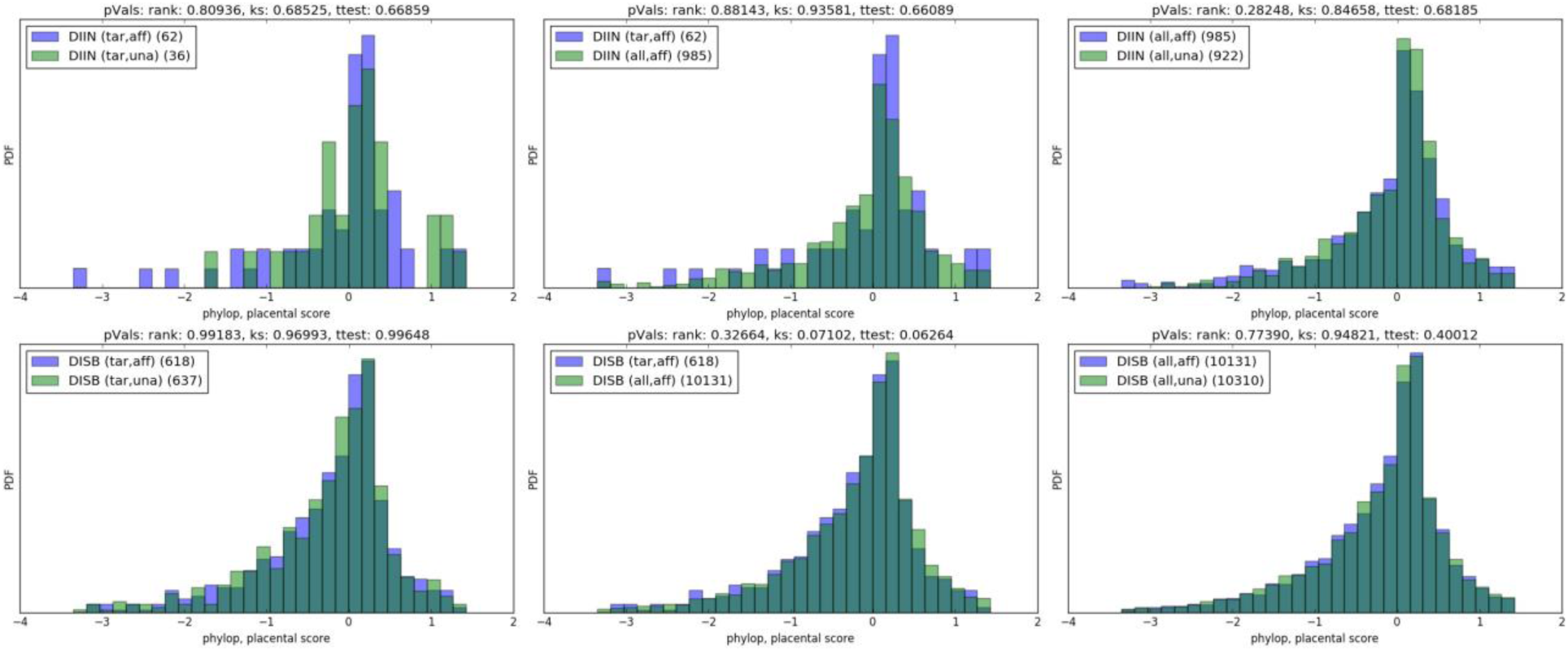
phylop, placental score distributions

**Figure S14.**
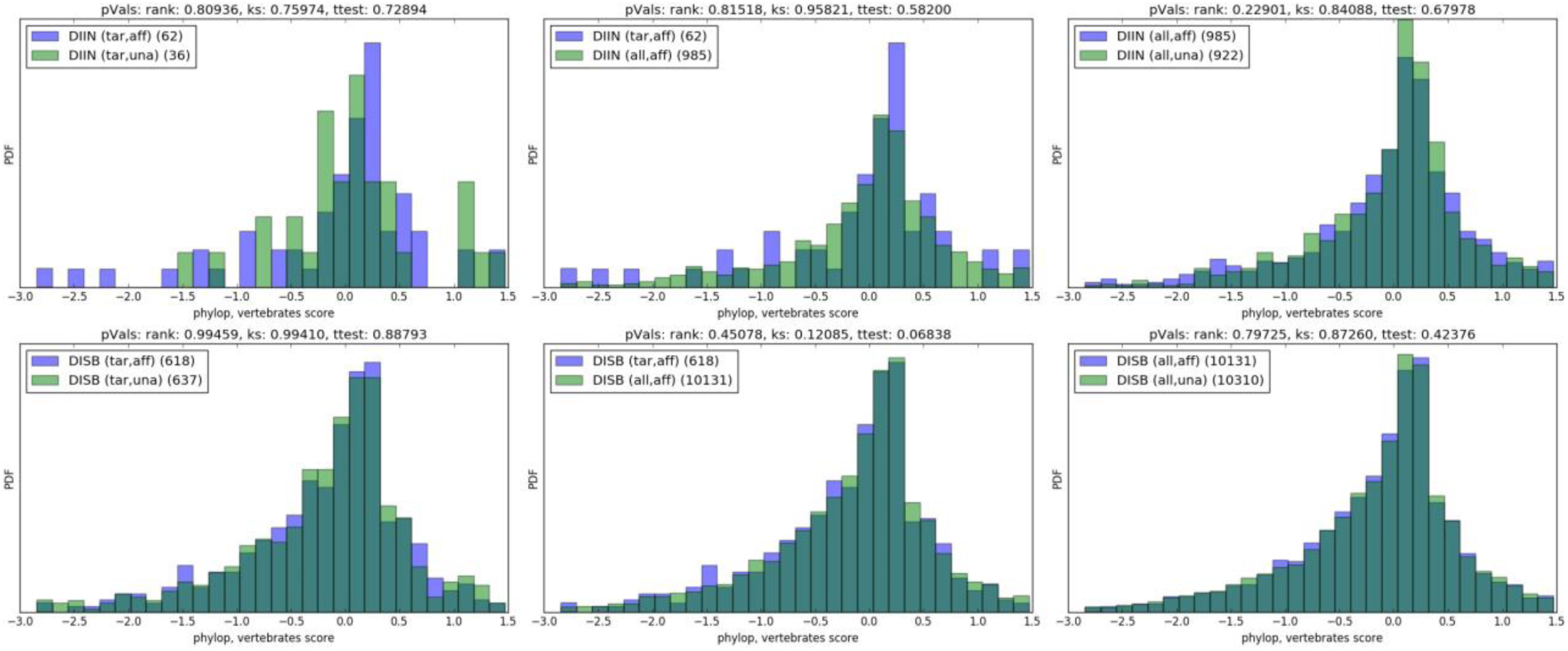
phylop, verbebrates score distributions

**Figure S15.**
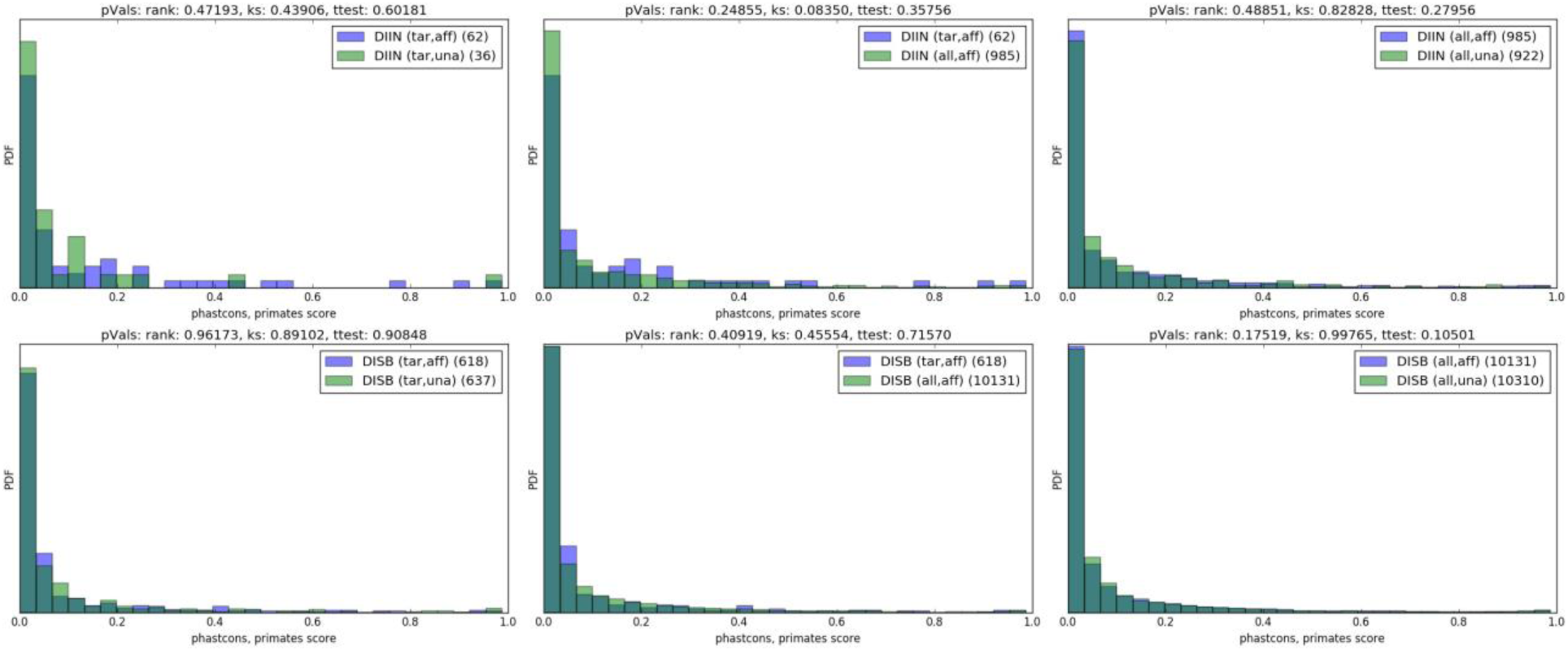
phastcons, primates score distributions

**Figure S16.**
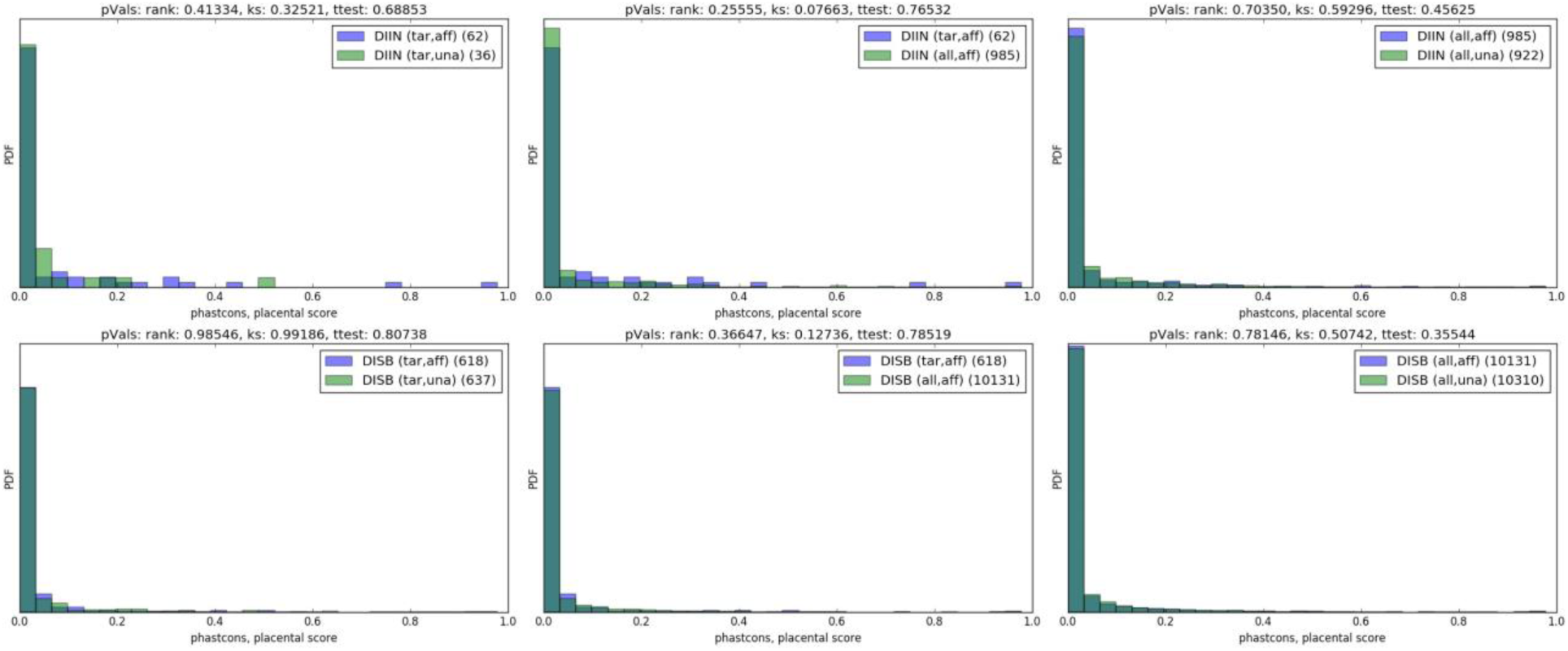
phastcons, placental score distributions

**Figure S17.**
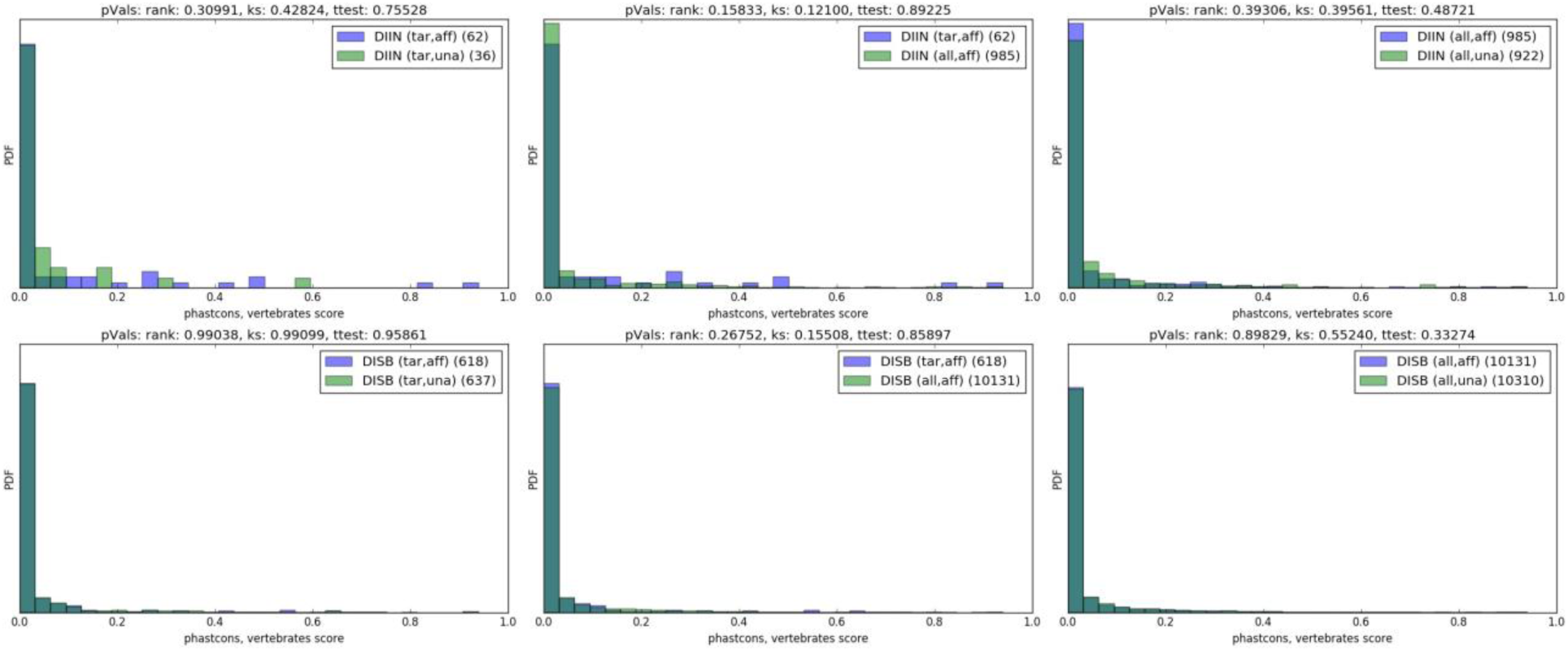
phastcons, vertabrates distributions

